# The transcription factor DksA exerts opposing effects on cell division depending on the presence of ppGpp

**DOI:** 10.1101/2023.05.15.540843

**Authors:** Sarah E. Anderson, Stephen E. Vadia, Jane McKelvy, Petra Anne Levin

## Abstract

Bacterial cell size is a multifactorial trait that is influenced by variables including nutritional availability and the timing of cell division. Prior work revealed a negative correlation between the alarmone (p)ppGpp (ppGpp) and cell length in *Escherichia coli*, suggesting that ppGpp may promote assembly of the division machinery (divisome) and cytokinesis in this organism. To clarify this counterintuitive connection between a starvation induced stress response effector and cell proliferation, we undertook a systematic analysis of growth and division in *E. coli* cells defective in ppGpp synthesis and/or engineered to overproduce the alarmone. Our data indicate that ppGpp acts indirectly on divisome assembly through its role as a global mediator of transcription. Loss of either ppGpp (ppGpp^0^) or the ppGpp-associated transcription factor DksA led to increased average length, with ppGpp^0^ mutants also exhibiting a high frequency of extremely long filamentous cells. Using heat-sensitive division mutants and fluorescently labeled division proteins, we confirmed that ppGpp and DksA are cell division activators. We found that ppGpp and DksA regulate division through their effects on transcription, although the lack of known division genes or regulators in available transcriptomics data strongly suggests that this regulation is indirect. Surprisingly, we also found that DksA inhibits division in ppGpp^0^ cells, contrary to its role in a wild-type background. We propose that the ability of ppGpp to switch DksA from a division inhibitor to a division activator helps tune cell length across different concentrations of ppGpp.

**Importance:** Cell division is a key step in the bacterial lifecycle that must be appropriately regulated to ensure survival. This work identifies the alarmone ppGpp as a general regulator of cell division, extending our understanding of the role of ppGpp beyond a signal for starvation and other stress. Even in nutrient replete conditions, basal levels of ppGpp are essential for division to occur appropriately and for cell size to be maintained. This study establishes ppGpp as a “switch” that controls whether the transcription factor DksA behaves as a division activator or inhibitor. This unexpected finding enhances our understanding of the complex regulatory mechanisms employed by bacteria to coordinate division with diverse aspects of cell growth and stress response. Because division is an essential process, a better understanding the mechanisms governing assembly and activation of the division machinery could contribute to the development of novel therapeutics to treat bacterial infections.

## Introduction

Bacteria must integrate information about their external environments and nutritional states to grow and replicate successfully. The alarmone nucleotides pppGpp and ppGpp [(p)ppGpp, hereafter collectively referred to as ppGpp] are widely conserved throughout the bacterial kingdom and serve as an intracellular signal for starvation and other stresses. In the model bacterium *Escherichia coli*, ppGpp levels are controlled by two RelA/SpoT Homologue (RSH)-family proteins: the synthetase RelA and the bifunctional synthetase/hydrolase SpoT (1). During amino acid starvation, binding of stalled ribosomes to RelA stimulates synthesis of ppGpp, raising its concentration as much as 100-fold over baseline in a phenomenon known as the stringent response (2, 3). Starvation for fatty acids, phosphate, carbon, or iron leads to SpoT-dependent accumulation of ppGpp, either by stimulating SpoT’s ppGpp synthesis activity (fatty acids, phosphate) or by inhibiting its ppGpp degradation ability (carbon) (4-6). Accumulation of ppGpp to high levels results in a reduction in overall biosynthesis, with exceptions for proteins required for adaptation to stressful conditions (1). ppGpp is also present at basal levels during balanced growth in nutrient replete conditions (7). Basal ppGpp is proposed to play a role in homeostasis (3), but a full understanding of the role of this molecule during balanced growth remains elusive.

In *E. coli* and other gamma-proteobacteria, ppGpp exerts its effects via transcriptional and post-translational mechanisms. As a transcriptional regulator, ppGpp binds to RNA polymerase (RNAP) at one of two equal-affinity sites, dubbed Sites 1 and 2 (8-10). Binding of ppGpp to RNAP during the stringent response leads to up- or downregulation of over 700 genes (11). Site 1 is comprised of RNAP subunits ω and β’ (9), while Site 2 is formed at the interface between RNAP and the transcription factor DksA (8). Unlike most transcription factors, DksA binds directly to RNAP to both activate and inhibit transcription (12, 13). DksA can bind RNAP with or without ppGpp, but binding of ppGpp at Site 2 leads to a conformational change in DksA that increases its affinity for RNAP (14). ppGpp cannot bind to Site 2 without DksA, and DksA and ppGpp do not appear to bind each other without RNAP (8). Usually, DksA and ppGpp work cooperatively to influence transcription; however, there are examples of ppGpp and DksA exerting independent or even opposing transcriptional and phenotypic effects (15-20). In addition to this transcriptional pathway, ppGpp also serves as a post-translational regulator by directly binding to at least 50 proteins to modulate their activities (3, 21-23). Together these two pathways allow ppGpp to modulate cell physiology and metabolism in response to environmental conditions and intracellular stress. Notably, although commonly thought of as a stress related alarmone, ppGpp is also important for homeostatic regulation during steady state growth, and defects in ppGpp synthesis lead to dysregulation of multiple pathways (7, 15, 18, 24-26).

To ensure survival, information about environmental conditions must be appropriately coordinated with the cell cycle, including the timing and frequency of cell division. Cell division is mediated by the divisome, a multiprotein complex (ten essential core proteins in *E. coli*) that guides the placement and synthesis of septal peptidoglycan at mid-cell (27). During growth, the members of the divisome assemble hierarchically at mid-cell. In *E. coli*, divisome assembly begins with the recruitment of the cytoplasmic tubulin homolog FtsZ to the future site of division. FtsZ serves as a dynamic scaffold for the rest of the complex. Divisome assembly terminates with FtsN (27). Divisome assembly is tightly controlled, with FtsZ serving as a major target of regulation during both balanced growth and stress (28-31). However, other components of the divisome can also be affected by regulation or environmental conditions; for example, FtsN is influenced by environmental pH (32). Regardless of mechanism, defects in division lead to increased cell length, while activation of division produces shorter cells.

Unsurprisingly, most components of the *E. coli* divisome are essential. In lieu of deletion mutations, conditional heat-sensitive alleles of division genes are extremely valuable for the study of divisome assembly and activation. Heat-sensitive mutations have been isolated in genes including *ftsZ*, *ftsA*, *ftsQ*, *ftsK*, and *ftsI* (33-36). When grown on LB without salt (LBNS), heat-sensitive mutants *ftsZ84*, *ftsA12*, and *ftsQ1* support growth at 30°C, but exhibit little to no growth at 37° or 42°C (32, 37). Mutants *ftsI23* and *ftsK44* have milder phenotypes, permitting growth at temperatures up to 37°C, but exhibiting reduced growth at 42°C on LBNS (32). Notably, the heat sensitivity of many of these alleles can be suppressed by gain-of-function mutations in or overexpression of other cell division genes (37-39), or by changes in the environment that promote divisome assembly (32). For example, the gain-of-function mutation *ftsA\** can suppress heat sensitivity of multiple conditional division mutants, including *ftsK44* and *ftsQ1* (40).

Significant data argue for a positive relationship between ppGpp and division. Initial evidence for divisome activation by ppGpp was reported in 1998 by Powell and Court, who found that ppGpp overproduction increased survival of a heat-sensitive *ftsZ* mutant (*ftsZ84*) at restrictive temperatures (41). In addition, there is a negative correlation between ppGpp levels and cell length (and to a more modest degree, cell width) (18, 25, 42, 43). Cells lacking ppGpp (referred to as ppGpp^0^) or DksA sometimes filament (18, 25). ppGpp^0^ cells are also more sensitive to increased production of SulA, an inhibitor of FtsZ assembly (26), suggesting that even basal levels of ppGpp have a positive impact on cell division. Both ppGpp^0^ cells and cells overexpressing ppGpp grow slowly despite their dramatic differences in length, strongly suggesting that ppGpp does not control size through its effects on growth rate (42). Modest increases in ppGpp cause resistance to mecillinam, an antibiotic targeting PBP2, the transpeptidase associated with the cell elongation machinery (elongasome) (44-46). Increased divisome activity can compensate for elongasome defects (47), suggesting that ppGpp causes mecillinam resistance by stimulating the divisome.

Substantial transcriptomics data argue against a model in which ppGpp regulates expression of *ftsZ* or any other division genes (11, 15, 44, 48, 49). Increases in ppGpp do not impact FtsZ protein levels (44), nor do FtsZ or other division proteins appear to be direct binding partners of ppGpp (21, 23) (J.D. Wang, personal communication). While the work of Powell and Court suggests that increases in ppGpp suppress the heat sensitivity of *ftsZ84* by increasing FtsZ84 production, they normalized FtsZ84 levels to levels of another protein NusA, which is negatively regulated by ppGpp (11, 15, 41, 49). This likely caused FtsZ84 levels to appear artificially elevated in their ppGpp overproducing strain. A more recent study reported a slight (less than two-fold) decrease in FtsZ levels normalized to OD_600_ in a ppGpp^0^ strain (26). The mechanism underlying this apparent decrease remains unclear, as it does not align with transcriptional data available for the ppGpp^0^ strain.

Many open questions remain about the mechanism by which ppGpp affects cell size and division. In particular, the contribution of basal ppGpp to cell size has not been carefully studied. To our knowledge the frequency of filamentation by ppGpp^0^ cells has never been quantified. Similarly, the effects of ppGpp on divisome assembly or on the survival of heat-sensitive division mutants other than *ftsZ84* have not been examined.

To gain insight into the mechanism by which ppGpp regulates cell division, we undertook a comprehensive analysis of the effect of ppGpp on divisome assembly and activation. We found that ppGpp is a positive regulator of divisome activity and assembly. Significantly, genetic interactions between ppGpp and *dksA* suggest that DksA exhibits opposing effects on division depending on the presence of ppGpp. Overall, this work suggests a nuanced mechanism allowing cell length to be adjusted to different concentrations of ppGpp.

## Results

### The impact of excess ppGpp on cell division is mediated through DksA

While previous work indicates that modest increases in ppGpp positively impact cell division in *E. coli* (41, 42), these studies did not determine the extent of this effect beyond suppression of *ftsZ84*, nor did they determine if ppGpp’s binding partner, DksA, was required. To address these gaps, we assessed the impact of excess ppGpp on *E. coli* cell size and division in the presence and absence of DksA. To increase ppGpp levels we took advantage of a plasmid (*prelA*), which encodes *relA* under an IPTG-inducible promoter (50, 51) (**Table 1, Table S1**). A control plasmid, *prelA’* encodes an inactive *relA* allele under the same promoter (**Table 1, Table S1**).

**Table 1.**
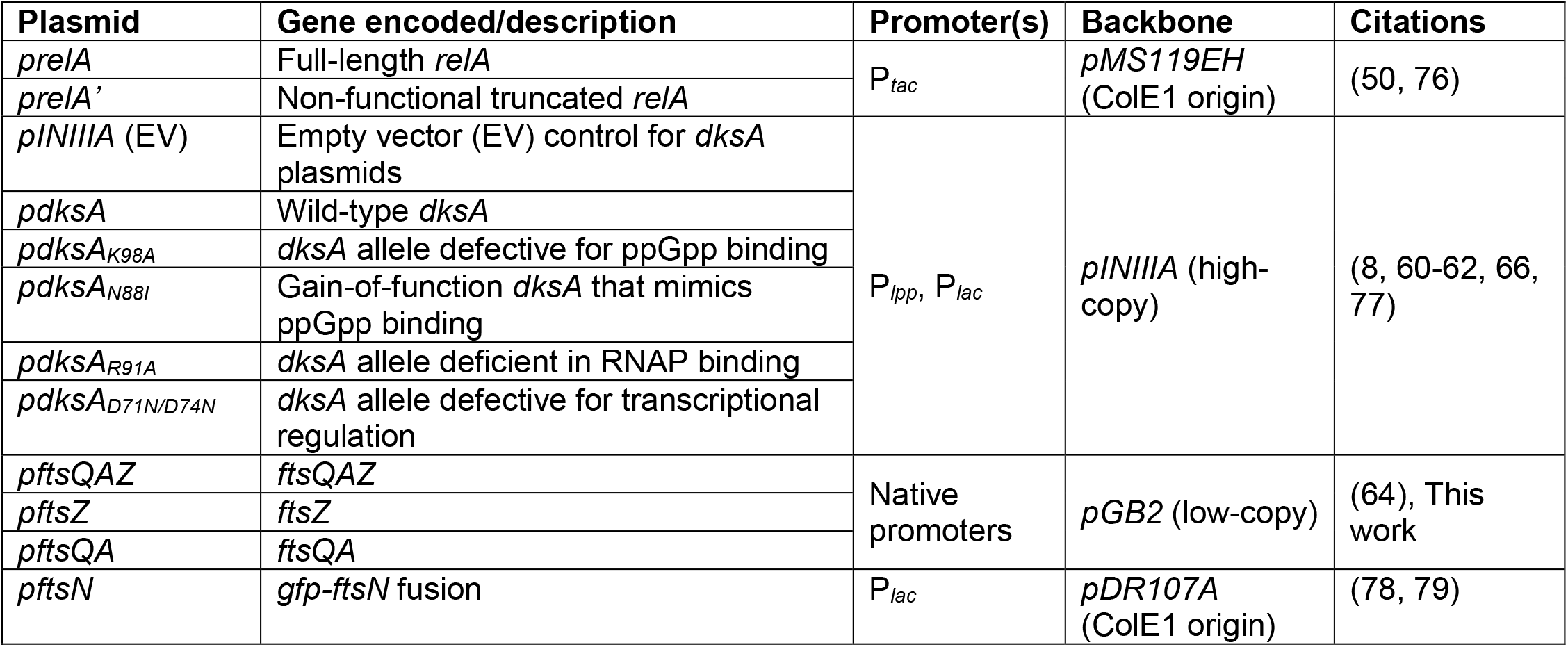
Expression plasmids used in this study. The gene(s) of interest encoded by each plasmid, the promoter(s) used to express them, and relevant information about each plasmid backbone are shown. Additional information about these plasmids, including their original published names and other encoded features, are available in Table S1.

As a first step we assessed the impact of excess ppGpp on cell size in the presence and absence of DksA. For these experiments we sampled cells at exponential phase (OD_600_ = 0.1), fixed, imaged, and measured for cell size using MicrobeJ (52). At least 200 cells were counted for each biological replicate. Notably, and in agreement with prior studies, the presence of *prelA* reduced *E. coli* cell size by ∼27% relative to *prelA’* in the absence of IPTG, suggesting read-through *relA* expression is sufficient to promote division (**Fig. 1A-D**) (42, 43). The difference in cell length mediated by excess ppGpp requires the presence of its RNAP binding partner, DksA. *prelA* resulted in only a ∼15% reduction in the length of *dksA*::*Kan* mutant cells, a difference not statistically significantly different from the *dksA*::*Kan prelA’* control (*P* = 0.056) (**Fig. 1E, F**). Importantly, the presence of *prelA* had no impact on cell width, consistent with a primary impact on division (**Fig. S1A**). The growth rates of *prelA* and *prelA’* strains were identical in the absence of inducer (**Fig. S1B, C**), indicating that under these conditions ppGpp mediates cell length independently of growth rate.

**Figure 1.**
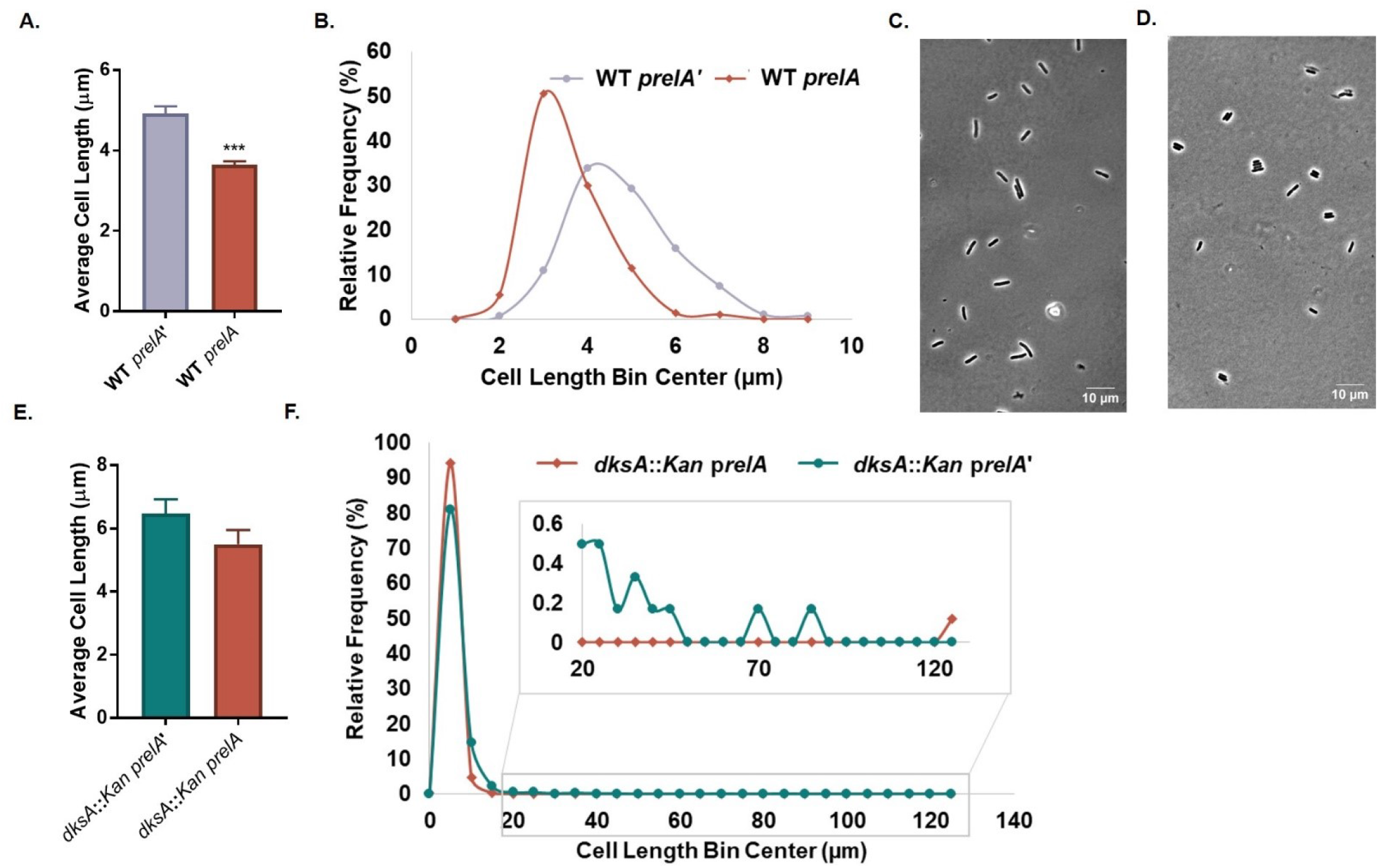
Excess ppGpp controls cell length through DksA. **A.** WT cells overexpressing *relA* (*prelA*) are shorter than cells overexpressing a truncated nonfunctional *relA* (*prelA’*). Data represent averages and SDs of three biological replicates (***, *P* ≤ 0.001, two-tailed t-test). **B.** Frequency distribution of lengths of individual cells expressing *prelA* or *prelA’*. N > 200 cells from a representative biological replicate (bin width = 1 µm). **C-D.** Representative phase contrast micrographs of WT *prelA’* (**C**) and WT *prelA* (**D**) cells. **E.** Cell length of *dksA*::*Kan* cells expressing *prelA* or *prelA’*. Data represent averages and SDs of three independent replicates (difference not significant by two-tailed t-test). **F.** Frequency distribution of individual cell lengths for *dksA*::*Kan* cells overexpressing *prelA* or *prelA’*. N > 600 cells from a single representative experiment (bin width = 5 µm).

To determine if DksA is required for ppGpp-mediated suppression of *ftsZ84*, as reported by Powell and Court (41), and to illuminate the impact of both DksA and ppGpp on the activity of other cell division proteins, we assessed the impact of excess ppGpp or *dksA* on the heat-sensitive alleles of five division genes: *ftsZ84* (G105S), *ftsA12* (A188V), *ftsQ1* (E125K), *ftsK44* (G80A), and *ftsI23* (Y380D) (34, 53-56). All five encode essential divisome components that assemble hierarchically at mid-cell (**Fig. 2A**) (27). For these experiments *relA* was ectopically expressed as above, while *dksA* was overexpressed from a plasmid containing both a constitutive and an IPTG-inducible promoter (**Table 1, Table S1**). To assess whether DksA is required for ppGpp’s documented effects on *ftsZ84* (41), we also expressed *prelA* in an *ftsZ84 dksA*::*Kan* double mutant. Strains were serially diluted and plated on LBNS without IPTG and incubated at 30°, 37°, and 42°C for 20 h prior to imaging.

**Figure 2.**
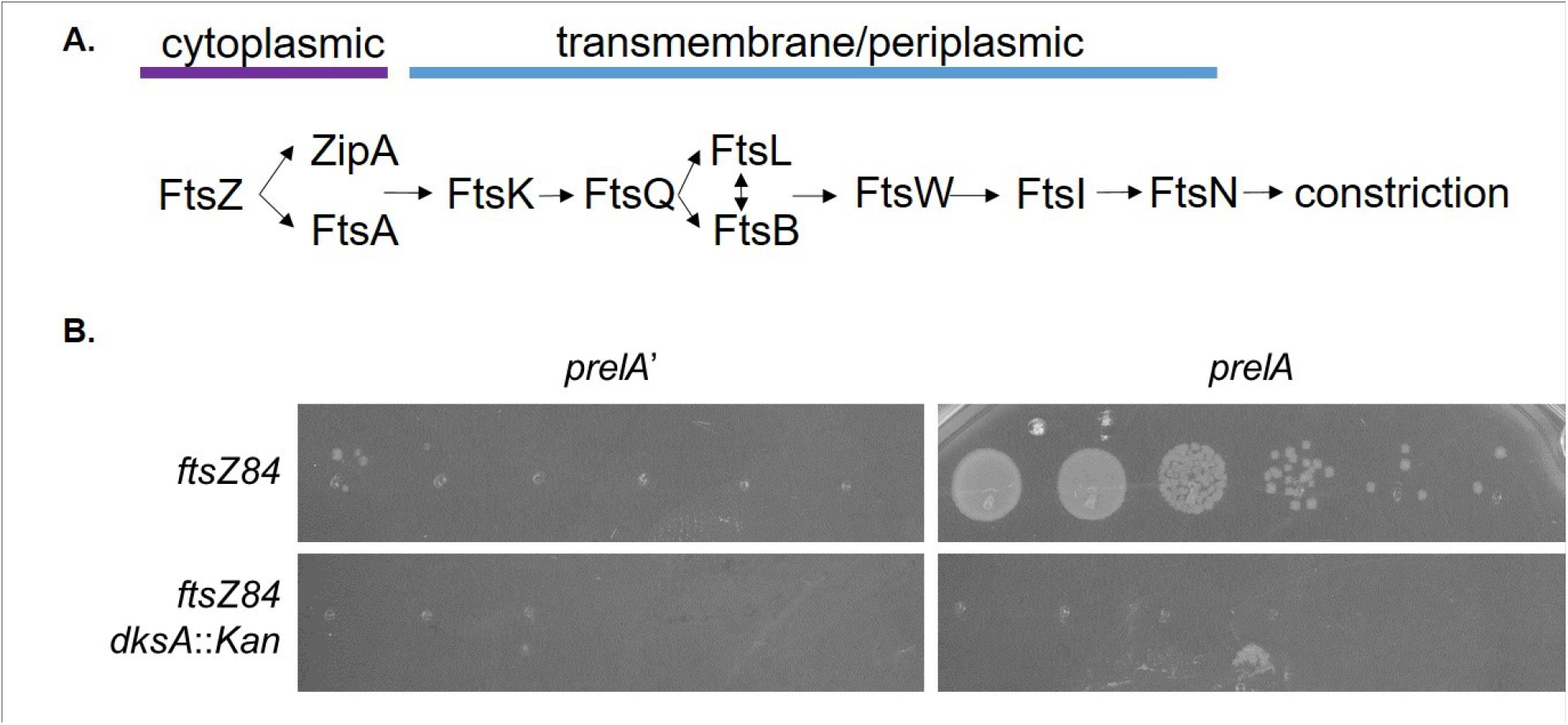
Excess ppGpp promotes division in a DksA-dependent manner. **A.** Essential division proteins assemble hierarchically to mid-cell. **B**. Expression of *prelA* facilitates growth of *ftsZ84*, but not *ftsZ84 dksA*::*Kan*, at 37°C. A representative image of three biological replicates is shown (no colonies are present for either *ftsZ84 dksA*::*Kan* strain).

Consistent with a role in ppGpp-mediated divisome activation, DksA was required for ppGpp-mediated suppression of *ftsZ84*. As expected based on previous work (41), *prelA* increased CFUs of *ftsZ84* by approximately four orders of magnitude at the restrictive temperature of 37°C (**Fig. 2B**). In contrast, an *ftsZ84 dksA*::*Kan prelA* strain was unable to form colonies at 37°C (**Fig. 2B**). Notably, *prelA* did not result in consistent changes in the heat sensitivity of other conditional division alleles (**Fig. S2A,**) suggesting that ppGpp and DksA influence division via FtsZ. Overexpression of *dksA* alone had no effect on division mutants, including *ftsZ84*, indicating that ppGpp is required for suppression of the latter (**Fig. S2B**).

### ppGpp facilitates divisome assembly and/or activation during steady-state growth

To clarify the contribution of ppGpp to division in nutrient replete conditions, we measured lengths of cells completely defective in ppGpp synthesis (Δ*relA spoT*::*cat*, hereafter ppGpp^0^) during growth in LB-glucose. As previously reported (18, 25), ppGpp^0^ mutants are heterogeneous for cell length; most ppGpp^0^ cells appeared slightly elongated compared to wild-type, while a sub-population of cells were highly filamentous (**Fig. 3A, B**). Considering the whole population, ppGpp^0^ cells were on average two-fold longer than the wild-type parent (**Fig. 3C**).

**Figure 3.**
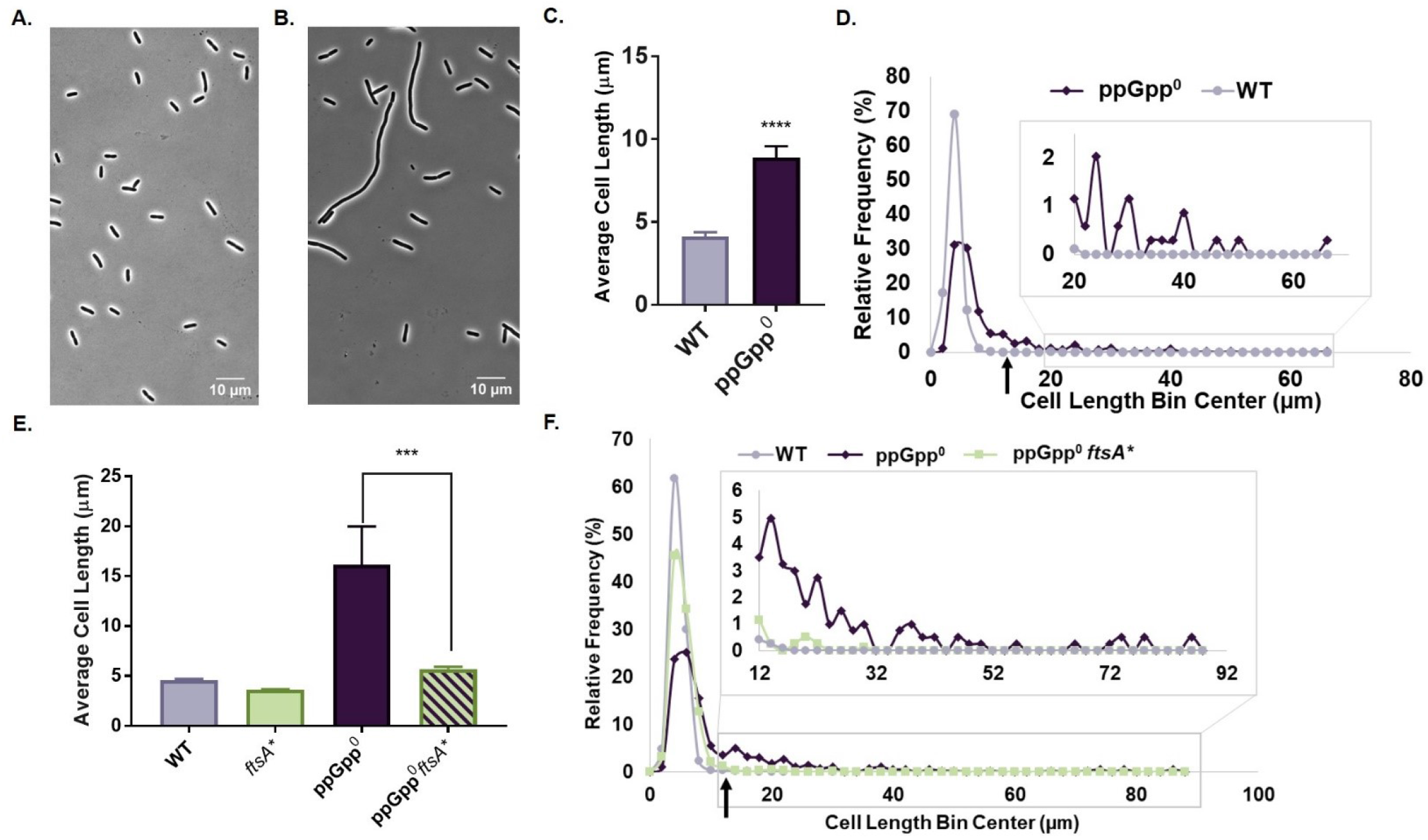
Basal ppGpp promotes cell division. **A-B.** Representative phase contrast micrographs of WT (**A**) and Δ*relA spoT*::*cat* (ppGpp^0^) (**B**) cells. **C.** ppGpp^0^ cells exhibit increased average cell length. Data represent averages and SDs of four biological replicates (****, *P* ≤ 0.0001, two-tailed t-test). **D.** Frequency distribution of individual cell lengths reveals both lengthening and filamentation among ppGpp^0^ cells. N > 200 cells from a representative biological replicate (bin width = 2 µm). The black arrow indicates the approximate location of the cut-off for filamentous cells (12.4 µm, 3x average wild-type length from Fig. 3C). **E.** Average cell length of ppGpp^0^ *ftsA\** cells is significantly reduced compared to the ppGpp^0^ parent strain. Data represent averages and SDs of at least three independent replicates (***, *P* ≤ 0.001 by one-way ANOVA with Tukey’s post-test). **F.** Frequency distribution shows decreased lengthening and filamentation in ppGpp^0^ *ftsA\** cells. N > 400 cells from a representative replicate (bin width = 2 µm). The black arrow indicates the approximate location of the cut-off for filamentous cells (13.4 µm, 3x average wild-type length from Fig. 3E).

To more fully characterize the impact of defects in ppGpp synthesis on cell length, we divided ppGpp^0^ cells into two populations using a cut-off of three times the average wild-type length to distinguish filaments from non-filaments (**Fig. 3D**). The non-filamentous subpopulation comprised ∼80% of the ppGpp^0^ cells (**Fig. 3D**). Notably, non-filaments were 46% longer than wild-type (average cell length ∼6.02 µm compared to 4.13 µm for wild-type), indicative of a significant defect in divisome assembly despite their relatively normal appearance. Filaments comprised ∼20% of the ppGpp^0^ population (**Fig. 3D**). The filamentous population had an average length of 22.06 µm, over five times longer than the wild-type average. Filamentous cells were exceedingly rare in the wild-type control, with cells exceeding 20 µm not observed (**Fig. 3D**). ppGpp^0^ cells had similar widths to wild-type under the conditions tested, reinforcing a primary defect in division (**Fig. S3A**). The ppGpp^0^ mutant grew 19% slower than wild-type in our conditions (**Fig. S3B)**.

To confirm that lengthening and filamentation of ppGpp^0^ are a consequence of reductions in divisome assembly and/or activity, we took advantage of a gain-of-function mutation in the cell division protein FtsA, FtsA*. FtsA*\** can compensate for a variety of defects in divisome assembly, including the heat-sensitive alleles *ftsQ1* and *ftsK44*, as well as complete loss of the normally essential proteins ZipA, FtsK, or FtsN (38-40, 57). FtsA* also exhibits increased interactions with FtsZ and increased recruitment of FtsN compared to wild-type FtsA (38, 58, 59). If the increased length of ppGpp^0^ cells is due to destabilization of the core divisome, then we would expect *ftsA\** to restore divisome function and decrease cell length and filamentation in this background.

Supporting a role for basal ppGpp in promoting divisome assembly at steady state, *ftsA\** reduced the average length of ppGpp^0^ (*relA*::*Kan spoT*::*cat*) cells ∼65% compared to the ppGpp^0^ parent strain (**Fig. 3E**). *ftsA\** also largely eliminated the heterogeneity of ppGpp^0^ mutants, reducing the fraction of filamentous cells from 33% to 2% (**Fig. 3F**). *ftsA\** also reduced the average length of non-filamentous ppGpp^0^ cells by 21%, from 6.77 µm to 5.37 µm. *ftsA\** did not significantly change the ppGpp^0^ growth rate (**Fig. S3C**).

Finally, we also measured the effect of loss of *dksA* on size. Defects in *dksA* had only a modest impact on cell length at steady-state in the presence of ppGpp, suggesting that the loss of the ppGpp-DksA-RNAP interaction is insufficient to explain the severe division defect in ppGpp^0^ cells. Non-filaments comprised ∼99% of the *dksA*::*Kan* population (**Fig. 4A-D**) (as compared to ∼80% of a ppGpp^0^ population (**Fig. 3D**)). On average *dksA*::*Kan* mutant cells were 17% longer than wild-type (**Fig. 4A**). Non-filamentous *dksA*::*Kan* cells had an average length of 4.97 µm, a 15% increase over wild-type, but 17% shorter than the non-filamentous ppGpp^0^ cells measured in **Fig. 3D**. *dksA*::*Kan* filaments (∼1% of the population) had an average length of 16.91 µm, a nearly four-fold increase compared to wild-type. *dksA*::*Kan* grew similarly to wild-type under the conditions tested (**Fig. S4**).

**Figure 4.**
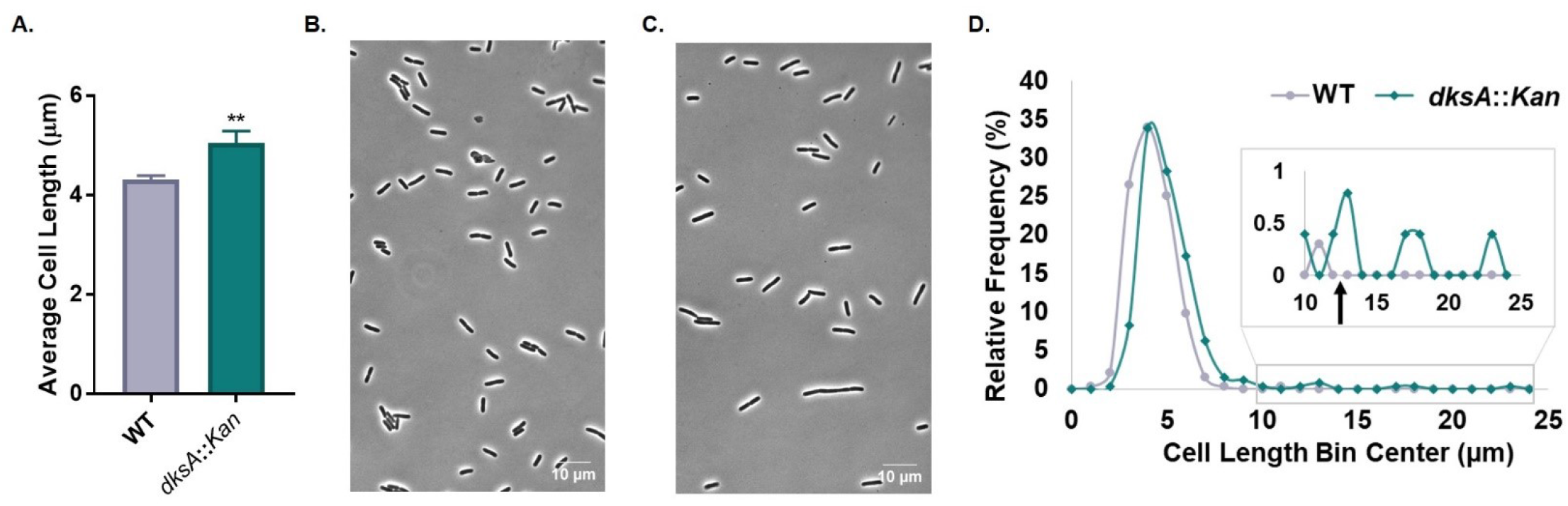
Deletion of *dksA* leads to cell lengthening. **A.** *dksA*::*Kan* cells exhibit increased average cell length relative to wild-type. Data represent averages and SDs of four biological replicates (**, *P* ≤ 0.01, two-tailed t-test). **B-C**. Representative phase contrast micrographs of WT (**B**) and *dksA*::*Kan* (**C**) cells. **D.** Frequency distribution of individual cells reveals lengthening and some filamentation by *dksA*::*Kan* cells compared to wild-type. N > 200 cells from a representative biological replicate (bin width = 1 µm). The black arrow indicates the approximate location of the cut-off for filamentous cells (12.9 µm, 3x wild-type cell length in Fig. 4A).

### ppGpp and DksA are required for optimal growth of conditional cell division mutants

Further supporting a role for basal ppGpp as well as DksA in divisome activation, we found *relA* and *dksA* to be required for optimal growth of multiple heat-sensitive mutants under permissive conditions (we were unable to obtain stable cell lines when we tried to combine heat-sensitive division alleles with mutations in both *relA* and *spoT*). Deletion of either *relA* (Δ*relA*, markerless deletion) or *dksA* (*dksA*::*Kan*) enhanced the heat sensitivity of *ftsZ84*, reducing CFUs at the permissive temperature of 30°C by roughly four orders of magnitude on LBNS (**Fig. 5**). Δ*relA* also reduced CFUs of *ftsK44* at the permissive temperature of 37°C by about three orders of magnitude, and decreased survival of *ftsA12* at both the permissive temperature of 30°C (by about two logs) and the non-permissive temperature of 37°C (no growth) (**Fig. 5**). Deletion of *relA* (*relA*::*Kan*) did not enhance heat killing of *ftsQ1* at the permissive temperature of 30°C; however, colony size was reduced (**Fig. 5**). *dksA*::*Kan* eliminated growth of both *ftsA12* and *ftsI23* at their non-permissive temperatures of 37°C and 42°C, respectively, and resulted in a roughly one log decrease in growth of *ftsK44* at the non-permissive temperature of 42°C (**Fig. 5**). *dksA*::*Kan* also decreased colony size and reduced CFUs of *ftsQ1* by about one order of magnitude at the permissive temperature of 30°C (**Fig. 5**). The effect of *relA*::*Kan* on *ftsI23* could not be determined due to the rapid accumulation of suppressors in the double mutant.

**Figure 5.**
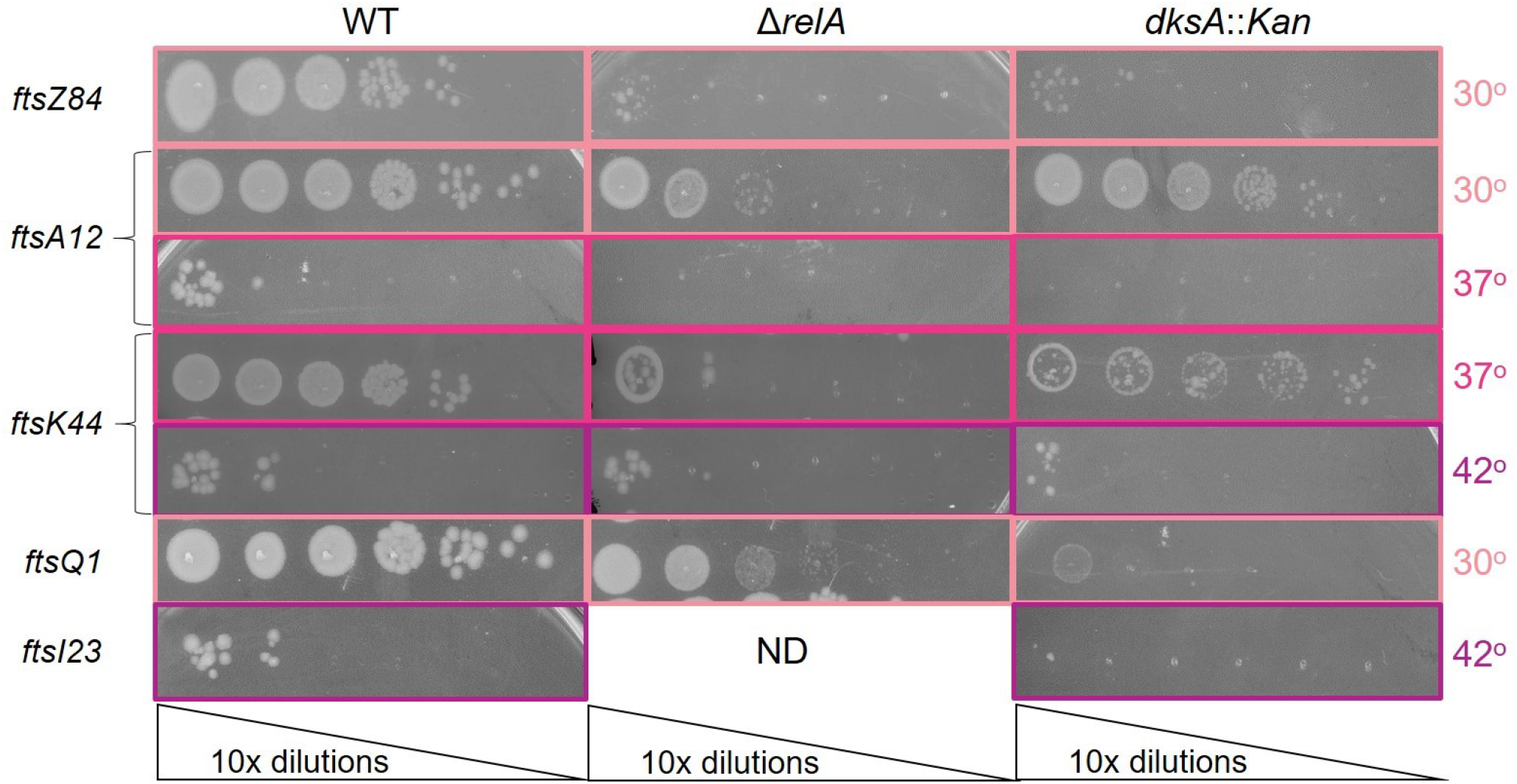
DksA and basal ppGpp are required for growth of conditional division mutants under permissive and non-permissive conditions. Effect of deletion of *relA* and *dksA* on heat-sensitive division mutants. Data shown are representative images of three biological replicates (ND, not determinable).

### DksA requires ppGpp to promote division

As noted in the introduction, interaction with ppGpp alters the DksA-RNAP interaction, and DksA displays different effects on transcription in the presence and absence of ppGpp (14, 15). To further illuminate the impact of ppGpp on DksA-mediated division effects, we took advantage of plasmids expressing separation-of-function *dksA* alleles affected for ppGpp binding (gift of R. Gourse) (**Table 1, Table S1, Fig. 6A**). These alleles include *dksA*_K98A_, which blocks binding of ppGpp to RNAP Site 2, and *dksA*_N88I_, which mimics ppGpp-bound DksA due to its enhanced affinity for RNAP (8, 60).

**Figure 6.**
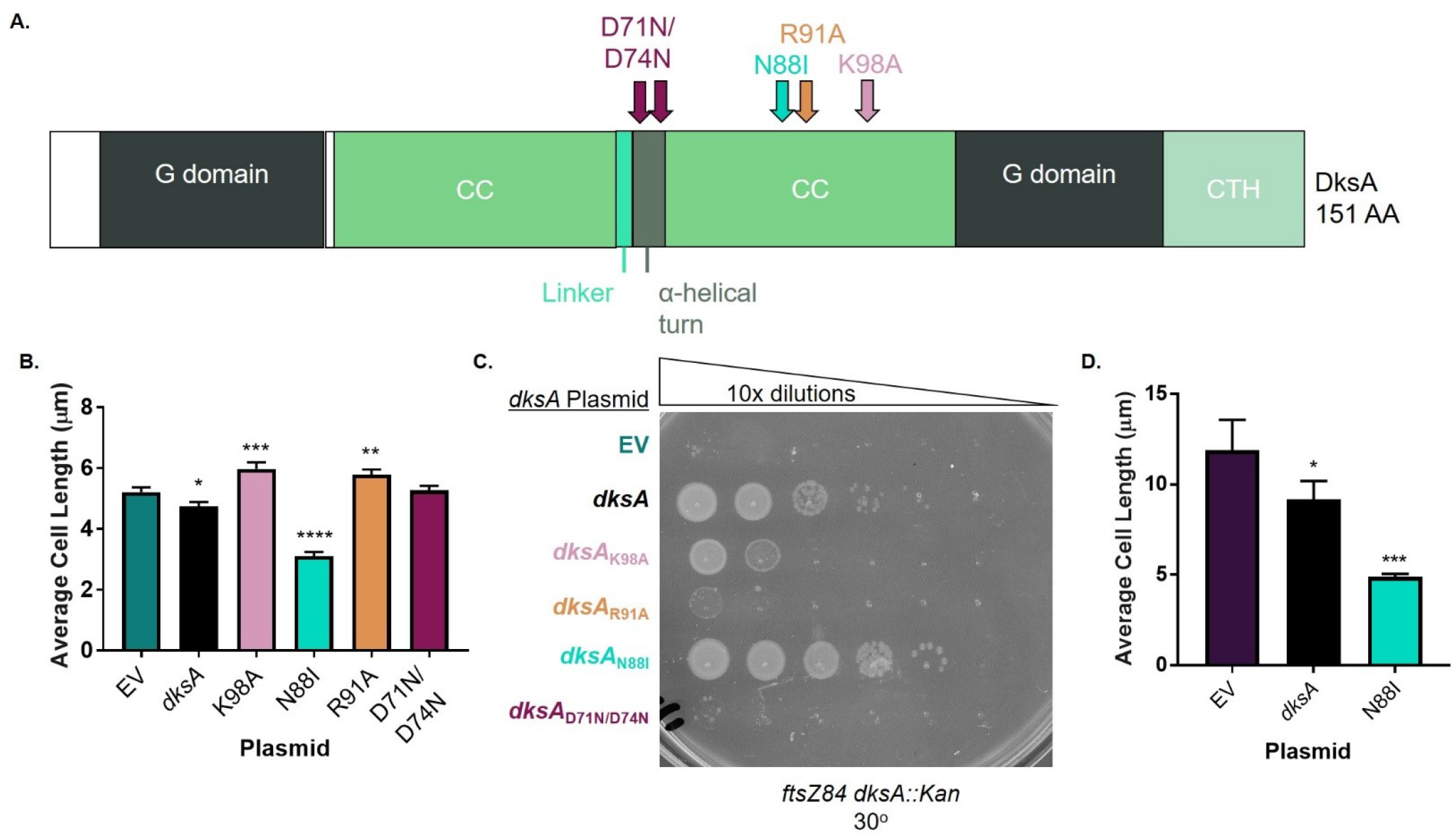
Mutant analysis indicates that DksA requires ppGpp and RNAP binding to promote division. **A.** Locations of separation-of-function mutations in DksA. DksA is comprised of a globular (G) domain, a coiled coil domain (CC) connected by a linker and α-helical turn, and a C-terminal helix (CTH) (80). In the α-helical turn, D71N/D74N interferes with transcriptional regulation by DksA. In the CC domain, N88I mimics ppGpp binding to DksA, R91A reduces DksA binding to RNAP, and K98A blocks binding of ppGpp to DksA. **B.** Complementation of cell length in a *dksA*::*Kan* mutant by different alleles of *dksA* expressed from a plasmid reveals that cell length control by DksA requires binding to ppGpp and RNAP. Data represent averages and SDs from three biological replicates (*, *P* ≤ 0.05; **, *P* ≤ 0.01; ***, *P* ≤ 0.001; ****, *P* ≤ 0.0001 relative to empty vector (EV) control by one-way ANOVA with Dunnett’s post-test). **C.** Complementation of enhanced *ftsZ84* heat sensitivity of a *dksA*::*Kan* mutant by different *dksA* alleles expressed from a plasmid supports a role for ppGpp and RNAP binding by DksA in the activation of FtsZ. A representative image of three biological replicates is shown. **D.** Effect of *dksA* alleles expressed from a plasmid on length of ppGpp^0^ cells shows that a gain-of-function *dksA* mutant can complement lengthening of ppGpp^0^ cells better than wild-type *dksA*. Data represent averages and SDs from three biological replicates (*, *P* ≤ 0.05; ***, *P* ≤ 0.001 relative to EV by one-way ANOVA with Dunnett’s post-test).

Analysis of cell length and *ftsZ84* heat sensitivity support a model in which ppGpp binding promotes DksA-dependent division activation. As expected, complementation of *dksA*::*Kan* with wild-type *dksA* significantly decreased length by 9% (**Fig. 6B, S5A**). The same plasmid also restored growth of *ftsZ84 dksA*::*Kan* at 30°C, increasing CFUs by three orders of magnitude (**Fig. 6C**). In contrast, a plasmid encoding the ppGpp “blind” *dksA* allele, *pdksA*_K98A_, failed to reduce the length of *dksA*::*Kan* cells (**Fig. 6B, S5A**), and resulted in only partial complementation of *ftsZ84 dksA*::*Kan*; CFUs were increased by about three logs, but colonies were extremely small at 30°C (**Fig. 6C**). Conversely, the DksA ppGpp-binding mimic, *dksA*_N88I_, decreased *dksA*::*Kan* cell length by ∼40%, even more than wild-type (**Fig. 6B, S5A**), and increased CFUs of *ftsZ84 dksA*::*Kan* by four orders of magnitude at 30°C (**Fig. 6C**). The growth rate of *dksA*::*Kan* mutants was unaffected by complementation with either the wild-type or mutant *dksA* alleles (**Fig. S5B**).

As a final confirmation that ppGpp-bound DksA is responsible for division activation, we expressed the ppGpp-binding mimic, *dksA*_N88I_, in ppGpp^0^ cells and measured the impact on length. *pdksA*_N88I_ reduced cell length of ppGpp^0^ by 58%, while *pdksA* reduced length by only 23% (**Fig. 6D**). *pdksA*_N88I_ reduced the percentage of filaments from 27% to ∼1%, while *pdksA* only reduced filamentation to 17%. (**Fig. S5C**). This suggests that the positive impact of ppGpp on division is mediated almost entirely through DksA. Both *pdksA* and *pdksA*_N88I_ increased growth rate of ppGpp^0^ (**Fig. S5D**). Altogether these experiments strongly indicate that ppGpp binding to Site 2 is required for activation of division by DksA.

### DksA functions as a division inhibitor in the absence of ppGpp

To gain a full picture of the effect of DksA and ppGpp on division, we measured length of ppGpp^0^ *dksA*::*Kan* triple mutant cells. To our surprise, our results suggested that DksA inhibits division in the absence of ppGpp. ppGpp^0^ *dksA*::*Kan* exhibited a 1.8-fold decrease in average cell length compared to ppGpp^0^ (**Fig. 7A**). This decrease in length was driven both by shortening of non-filamentous cells and by a decrease in the number of filaments. Non-filamentous ppGpp^0^ *dksA*::*Kan* cells had an average length of 4.81 µm, compared to 5.67 µm for non-filamentous ppGpp^0^ and 4.03 µm for wild-type. The frequency of filaments was reduced from 21% in ppGpp^0^ to 3% in ppGpp^0^ *dksA*::*Kan* (**Fig. 7B-E**). Altogether, ppGpp^0^ *dksA*::*Kan* looked more similar to *dksA*::*Kan* than ppGpp^0^, suggesting that DksA may be epistatic to ppGpp in terms of cell length (**Fig. 4**). The ppGpp^0^ *dksA*::*Kan* mutant also exhibited a faster growth rate than the ppGpp^0^ parent (**Fig. S6**).

**Figure 7.**
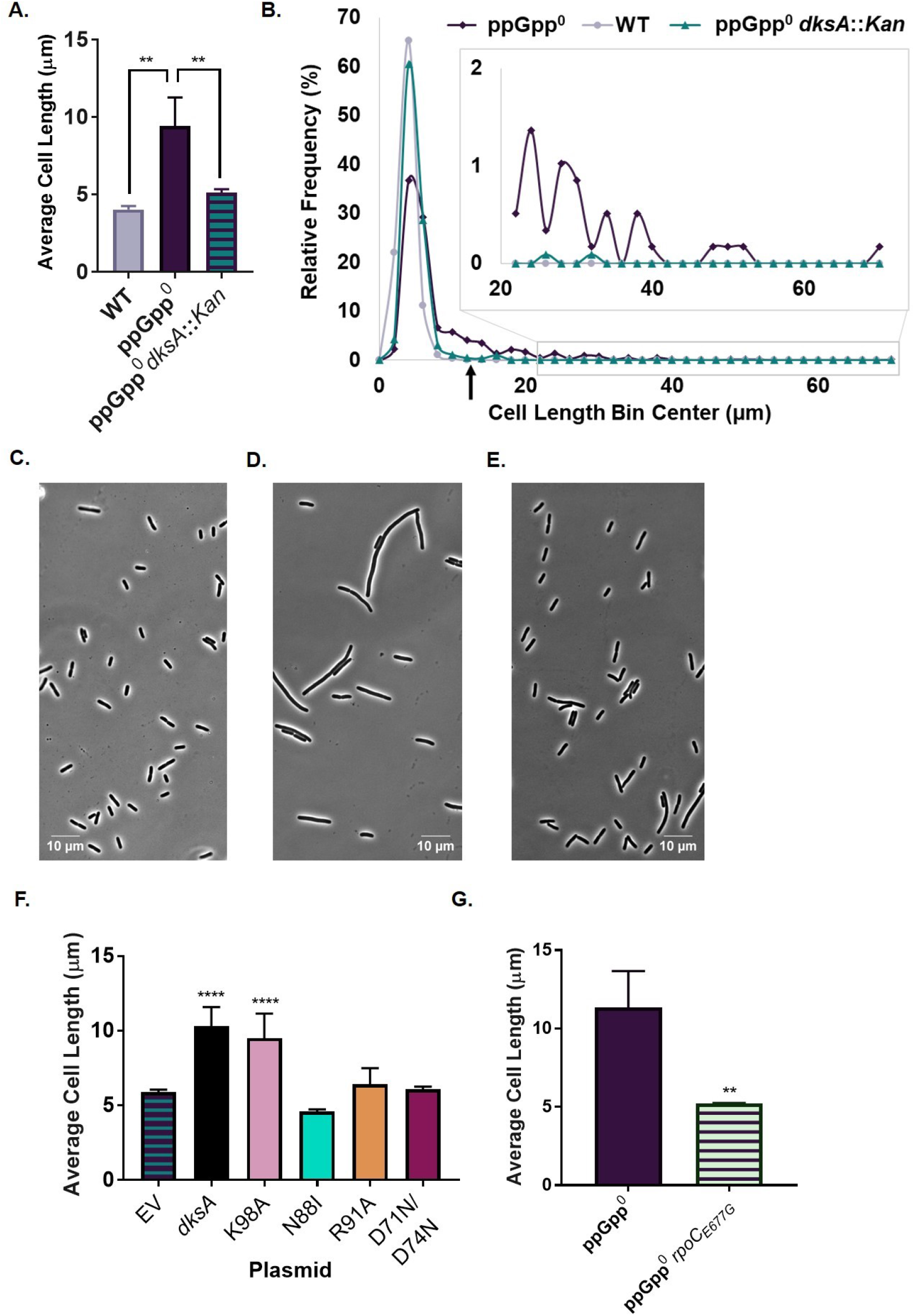
In the absence of ppGpp, DksA behaves as a division inhibitor. **A.** Deletion of *dksA* in the ppGpp^0^ mutant reduces average cell length. Data represent averages and SDs of three independent replicates (**, *P* ≤ 0.01 relative to ppGpp^0^ by one-way ANOVA with Dunnett’s post-test). **B.** Frequency distribution of single cell lengths demonstrates that ppGpp^0^ *dksA*::*Kan* exhibits less lengthening and filamentation than ppGpp^0^, although cells are still longer and exhibit more filaments than wild-type. N > 500 cells from a representative experiment (bin width = 2 µm). The black arrow indicates the approximate location of the cut-off for filamentous cells (12.2 µm, 3x wild-type length in Fig. 7A). **C-E.** Representative phase contrast micrographs of WT (**C**), ppGpp^0^ (**D**), and ppGpp^0^ *dksA*::*Kan* (**E**) cells. **F.** Complementation of decreased length in ppGpp^0^ *dksA*::*Kan* mutants by different alleles of *dksA* expressed from plasmids reveals a requirement for RNAP binding by DksA. Data represent averages and SDs of at least three biological replicates (****, *P* ≤ 0.0001 relative to empty vector control by one-way ANOVA with Dunnett’s post-test). **G.** An *rpoC* mutation that blocks binding of DksA to RNAP decreases average length in a ppGpp^0^ background. Data represent averages and SDs of three biological replicates (**, *P* ≤ 0.01 by two-tailed t-test).

To further confirm that DksA acts as a division inhibitor when not bound to ppGpp, we complemented ppGpp^0^ *dksA*::*Kan* with plasmids encoding wild-type *dksA*, the ppGpp blind allele, *dksA*_K98A_, and the ppGpp bound mimic, *dksA*_N88I_, and measured effects on cell length. As expected, a plasmid encoding wild-type *dksA* increased cell length of ppGpp^0^ *dksA*::*Kan* 1.8-fold and increased the frequency of filamentation from 3% to 18% (**Fig. 7F, S7A**). Complementation with the wild-type allele also reduced the growth rate of ppGpp^0^ *dksA*::*Kan* (**Fig. S7B**). Consistent with DksA inhibiting division when not bound to ppGpp, complementation of ppGpp^0^ *dksA*::*Kan* with the ppGpp blind allele, *pdksA*_K98A_, increased cell length by 1.6-fold and increased the frequency of filaments to 16% (**Fig. 7F, S7B**). In contrast, the ppGpp binding mimic, *pdksA*_N88I_ failed to complement cell length and actually reduced filamentation to ∼0.5% (**Fig. 7F, S7B**), supporting a model in which ppGpp binding switches DksA from a division inhibitor to a division activator.

### DksA and ppGpp indirectly regulate the divisome through transcription

Although substantial transcriptomics data argue against either ppGpp or DksA as direct regulators of division gene expression, the most straightforward explanation of their positive influence on division is via interaction with RNAP. To probe this idea, we complemented *dksA*::*Kan*, *dksA*::*Kan ftsZ84*, and ppGpp^0^ *dksA*::*Kan* mutants with plasmids encoding *dksA* alleles defective for interactions with RNAP (**Table 1, Table S1**). These alleles include: *dksA*_R91A_, which has reduced affinity for RNAP, and *dksA*_D71N/D74N_, which still binds RNAP but is defective for regulating transcriptional initiation by RNAP (8, 61, 62) (**Fig. 6A**).

Altogether our results support a model in which DksA and ppGpp regulate division through interaction with RNAP and downstream effects on transcription. In contrast to wild-type *dksA*, both *dksA*_R91A_ and *dksA*_D71N/D74N_ failed to complement the cell length associated with *dksA*::*Kan* (**Fig. 6B, S5A**) or facilitate growth of *ftsZ84 dksA*::*Kan* at 30°C (**Fig. 6C**). *pdksA*_R91A_ and *pdksA*_D71N/D74N_ also failed to increase length or the rate of filamentation of ppGpp^0^ *dksA*::*Kan* (**Fig. 7F, S7B**).

To further confirm whether DksA binding to RNAP is required for inhibition of division in the absence of ppGpp, we measured cell length of a ppGpp^0^ *rpoC*_E677G_ mutant. The *rpoC*_E677G_ mutation blocks binding of DksA to RNAP (63). This mutation reduced cell length in the ppGpp^0^ background by approximately two-fold (**Fig. 7G**) and reduced the percentage of filamentous cells from 24% to ∼1% (**Fig. S9A**), although the growth rate was not affected (**Fig. S9B**). This strongly suggests that DksA’s effects on transcription by RNAP are responsible for inhibition of division in the absence of ppGpp.

### A ppGpp binding defective RNAP_1-2-_ mutant does not phenocopy length and division phenotypes associated with loss of ppGpp

In an attempt to further confirm that ppGpp activates division through transcription, we measured cell length and *ftsZ84* heat sensitivity in a strain containing mutations in *rpoZ* and *rpoC* that block binding of ppGpp to RNAP (designated RNAP_1-2-_) (8). Unexpectedly, RNAP_1-2-_ mutants failed to phenocopy a ppGpp^0^ strain, exhibiting no change in length or filamentation compared to an isogenic wild-type control (RNAP_1+2+_) (**Fig. S8A, B**). A RNAP_1-2-_ *ftsZ84* mutant exhibited a roughly one log decrease in CFUs at the permissive temperature of 30°C (**Fig. S8C**), a much milder change than that caused by Δ*relA* (**Fig. 5**). Furthermore, the RNAP_1-2-_ *ftsZ84* strain actually exhibited a two log increase in CFU at the non-permissive temperature of 37°C, again in contrast to *ftsZ84* Δ*relA* (**Fig. S8C, D**). Available transcriptomics data do not provide an obvious explanation for this unexpected discrepancy (11).

### ppGpp and DksA promote recruitment of divisome proteins to the nascent septum

Altogether our data suggest ppGpp and DksA work coordinately to modulate cell division during both exponential growth and nutrient stress. To illuminate the step at which they function, we measured recruitment of fluorescently labeled division proteins to the nascent septum in wild-type, ppGpp^0^, *dksA*::*Kan*, and ppGpp^0^ *dksA*::*Kan* strains. In wild-type cells, the divisome complex forms a ring-like structure at mid-cell, which can be visualized as a bright band of fluorescence using GFP-labeled proteins (**Fig. S10A-E**). A defect in recruitment for a given protein fusion should result in a reduction of the fraction of cells exhibiting a bright band at mid-cell.

For these experiments we expressed *gfp-*tagged, IPTG-inducible alleles of the essential cell division genes *ftsZ*, *ftsA*, *ftsL*, *ftsI*, and *ftsN* from the lambda locus in strains also encoding a wild-type copy of that gene, as described previously (32). Strains were grown in LB IPTG and fixed prior to imaging. To quantify changes in division protein recruitment between strains, we calculated the “length-to-ring ratio” (L/R), which is the total cell length for the entire population divided by the total number of rings. An increase in L/R indicates a defect in division protein recruitment.

Supporting DksA as a division inhibitor in the absence of ppGpp, ppGpp^0^ cells exhibit DksA-dependent defects in divisome recruitment. ppGpp^0^ exhibited significant 2-3-fold increases in L/R for all division proteins tested, indicating defects in divisome assembly (**Fig. 8A**). Filamentous ppGpp^0^ cells usually had only one or two rings of each division protein, and those rings were typically located near the ends of filaments (**Fig. S10**). *dksA*::*Kan* exhibited no changes in L/R for any division proteins tested compared to wild-type (**Fig. 8A**). On the other hand, ppGpp^0^ *dksA*::*Kan* exhibited significant ∼2-fold reductions in L/R of GFP-FtsZ, GFP-FtsL, and GFP-FtsI compared to ppGpp^0^ (**Fig. 8B**). Similar trends were observed for GFP-FtsA and GFP-FtsN, although differences were not statistically significant. For all division proteins, L/R was indistinguishable between *dksA*::*Kan* and ppGpp^0^ *dksA*::*Kan*, indicating that DksA is epistatic to ppGpp in terms of division protein recruitment. These results suggest that in the absence of ppGpp, DksA inhibits divisome assembly, but ppGpp activates assembly by binding DksA and relieving this inhibition.

**Figure 8.**
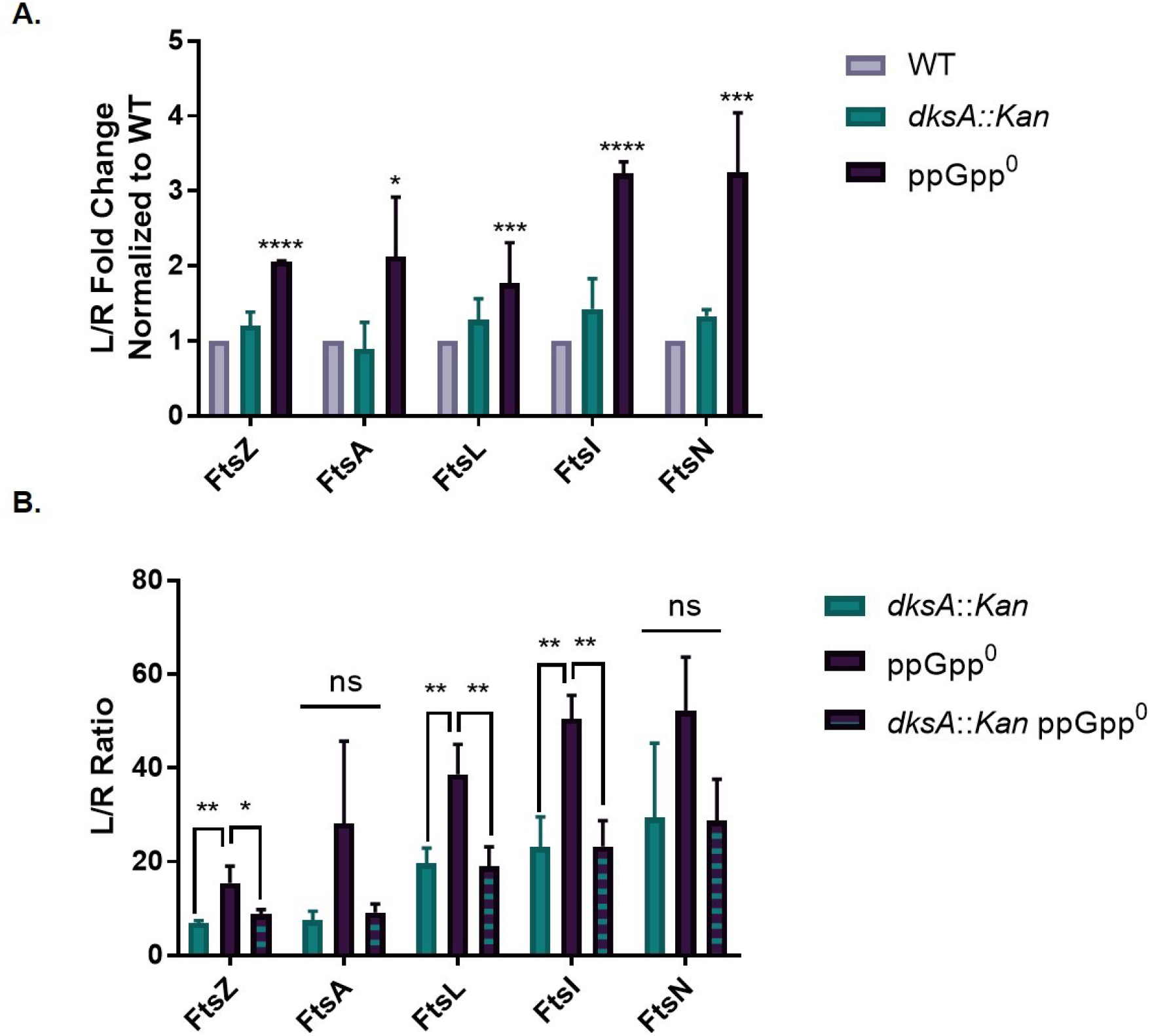
ppGpp and DksA modulate division protein recruitment. **A.** Fold changes of length/ring ratios for GFP-tagged division proteins expressed in *dksA*::*Kan* and ppGpp^0^ cells relative to wild-type. Data represent averages and SDs of three independent replicates (*, *P* ≤ 0.05; ***, *P* ≤ 0.001; ****, *P* ≤ 0.0001 relative to WT by one-way ANOVA of untransformed data with Dunnett’s post-test). **B.** Length/ring ratios of GFP-tagged division proteins expressed in *dksA*::*Kan*, ppGpp^0^, and ppGpp^0^ *dksA*::*Kan* cells. Data represent averages and SDs of three biological replicates (ns, not significant; *, *P* ≤ 0.05; **, *P* ≤ 0.01 by one-way ANOVA with Tukey’s multiple comparisons test).

### DksA and ppGpp likely modulate divisome assembly through their impact on recruitment of FtsZ and FtsN

Thus far, our data support a model in which transcriptional changes mediated by ppGpp and DksA lead to indirect activation (DksA + ppGpp) or inhibition (DksA alone) of divisome assembly. At the same time, the hierarchical nature of divisome assembly makes it difficult to know which proteins are the primary target. To clarify this issue, we examined the effect of overexpressing a collection of division genes on cell length in *dksA*::*Kan* and ppGpp^0^ backgrounds.

Supporting FtsZ as a primary target, the presence of a plasmid encoding *ftsZ* under the control of its native promoter was sufficient to reduce the length of both *dksA*::*Kan* and ppGpp^0^ cells. Expression of the *ftsQAZ* operon from its native promoter on a low copy plasmid (64) (**Table 1, Table S1**) reduced cell length of ppGpp^0^ by 2.8-fold and reduced length of *dksA*::*Kan* by 37% (**Fig. 9A**). *pftsQAZ* also reduced the proportion of filaments in the ppGpp^0^ background from 30% to 3% (**Fig. 9B**). Similar results were obtained with a plasmid overexpressing only *ftsZ*, which reduced length of ppGpp^0^ and *dksA*::*Kan* by 2.8-fold and 17%, respectively, and reduced filamentation of ppGpp^0^ from 37% to 3% (**Fig. 9C, D**). A plasmid overexpressing only *ftsQA* failed to significantly affect size in either background, indicating these genes are unlikely to be directly involved in ppGpp-mediated division activation (**Fig. S11A**). *pftsQAZ* and *pftsZ* also reduced wild-type cell length by 30% and 11%, respectively (**Fig. 9A, B**).

**Figure 9.**
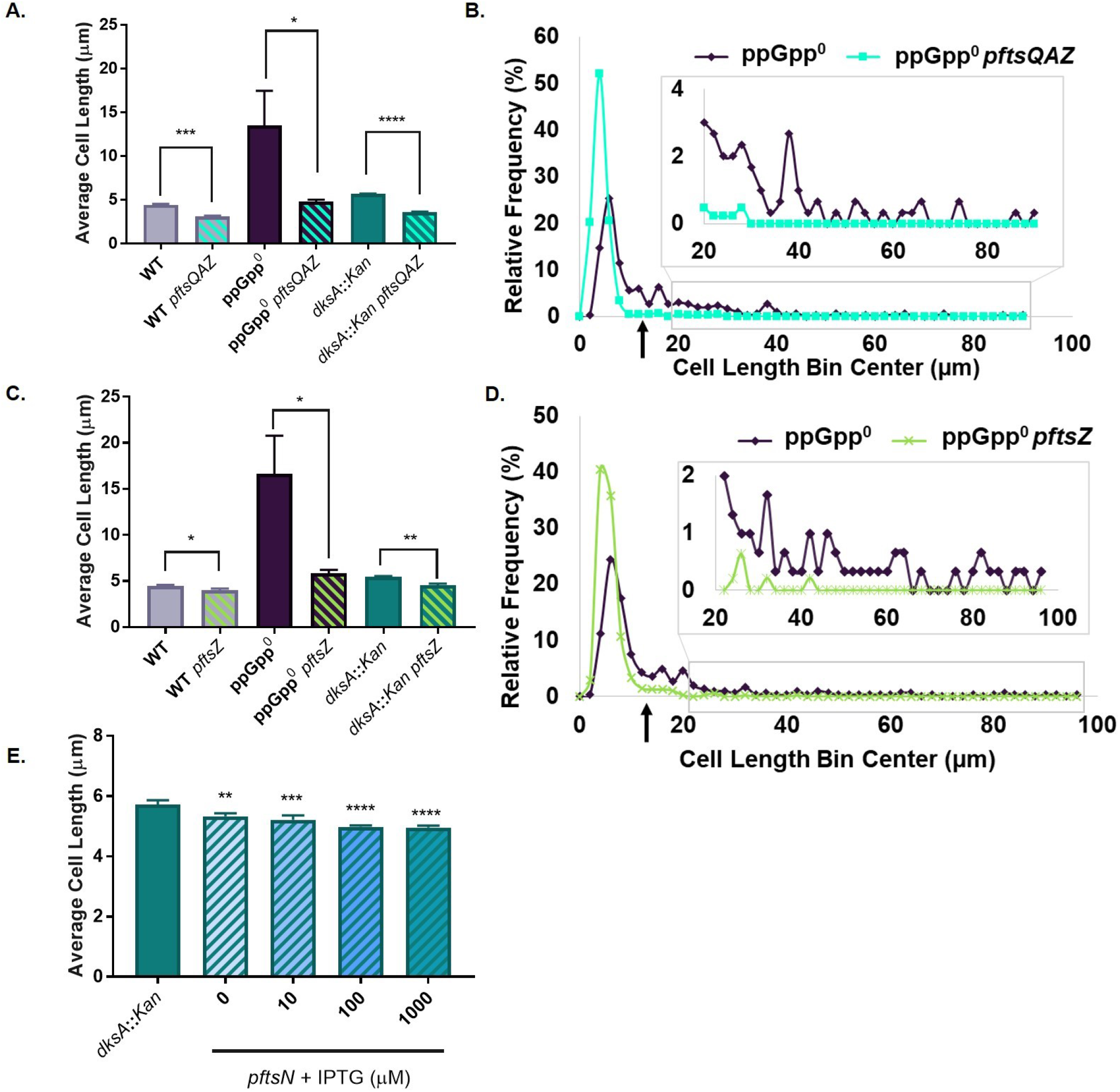
Overexpression of *ftsZ* or *ftsN* reduces length of ppGpp^0^ (*ftsZ* only) and *dksA*::*Kan* (both). **A.** Overexpression of the *ftsQAZ* operon leads to decreased average cell length in ppGpp^0^ and *dksA*::*Kan* cells relative to vector-free controls. Data represent averages and SDs of three biological replicates (**, *P* ≤ 0.01; ****, *P* ≤ 0.0001 by one-way ANOVA with Tukey’s post-test). **B.** Frequency distribution of individual cell lengths shows that overexpression of *ftsQAZ* decreases both lengthening and filamentation among ppGpp^0^ cells. N > 200 cells from a single representative experiment (bin width = 2 µm). The black arrow indicates the approximate cut-off for filamentous cells (13.4 µm, 3x the wild-type average in Fig. 9A). **C.** Overexpression of *ftsZ* alone leads to decreased average length of wild-type, ppGpp^0^, and *dksA*::*Kan* cells relative to vector-free controls. Data represent averages and SDs of three biological replicates (*, *P* ≤ 0.05; **, *P* ≤ 0.01 by two-tailed t-test). **D.** Frequency distribution of individual cell lengths shows that overexpression of *ftsZ* alone reduces lengthening and filamentation among ppGpp^0^ cells. N > 300 cells from a single representative experiment (bin width = 2 µm). The black arrow indicates the approximate cut-off for filamentous cells (13.5 µm, 3x the wild-type average in Fig. 9C). **E.** Overexpression of *gfp-ftsN* from an IPTG-inducible plasmid leads to significant decreases in cell length in the *dksA*::*Kan* background. Data represent averages and SDs of three biological replicates (**, *P* ≤ 0.01; ***, *P* ≤ 0.001; ****, *P* ≤ 0.0001 relative to *dksA*::*Kan* by one-way ANOVA with Dunnett’s post-test).

Intriguingly, overexpression of *ftsN,* encoding the last protein recruited to the division site, the so-called “trigger” FtsN, reduced the length of *dksA*::*Kan* mutants but not ppGpp^0^ cells, suggesting that DksA may mediate division through its impact on both FtsZ and FtsN. Expression of *gfp-ftsN* from an IPTG-inducible promoter reduces the length of wild-type cells (32), and also reduced length of *dksA*::*Kan* cells by up to 13% at the highest level of induction (**Fig. 9E**). *gfp-ftsN* expression, however, had no impact on ppGpp^0^ cell length (**Fig. S11B**). No changes in growth rate were observed for strains expressing *pftsQAZ, pftsZ*, or *pftsN* (**Fig. S12A-D**).

### ppGpp and DksA modulate division independently of FtsZ concentration

Because different groups have reported different effects of ppGpp on FtsZ protein levels (26, 41, 44), and our results suggest that FtsZ is a major target of regulation by DksA and ppGpp, we wanted to clarify whether DksA and ppGpp regulate FtsZ levels via quantitative immunoblotting. We quantified FtsZ concentrations in wild-type, ppGpp^0^, *dksA*::*Kan*, and ppGpp^0^ *dksA*::*Kan*, as well as in wild-type cells expressing *prelA, prelA’,* or *pftsQAZ*. We found that DksA and ppGpp have no effect on FtsZ levels. Only the *pftsQAZ* positive control exhibited a significant change in FtsZ concentration compared to wild-type (**Fig. S13A-D**). This demonstrates that although ppGpp and DksA influence divisome assembly through FtsZ, they do not regulate FtsZ levels.

## Discussion

This study establishes ppGpp and DksA as critical modulators of cell length and division in *E. coli*. Most importantly, when bound to ppGpp, DksA interacts with RNAP to promote assembly of the division machinery and reduce cell length (**Fig. 10A**). Loss of *dksA* alone leads to increased cell length (**Fig. 4, 10C**). While extensive transcriptomics data suggest ppGpp-DksA’s impact on division is indirect and independent of changes in division gene expression (11, 15, 44, 48, 49), genetic data suggest FtsZ and FtsN are the eventual downstream targets (**Fig. 9**). Unexpectedly, in the absence of ppGpp, DksA has an inhibitory effect on division (**Fig. 10B**). Deletion of *dksA* in the absence of ppGpp significantly reduces the fraction of filamentous cells in the population and also reduces the average length of the non-filamentous fraction (**Fig. 7, 10D**). Genetic and cytological analysis suggest that DksA exerts an inhibitory effect on division via changes in recruitment and/or activation of FtsZ (**Fig. 9**). Transcriptomic data indicate that the *negative* impact of DksA on division in the absence of ppGpp is also likely indirect (15, 48).

**Figure 10.**
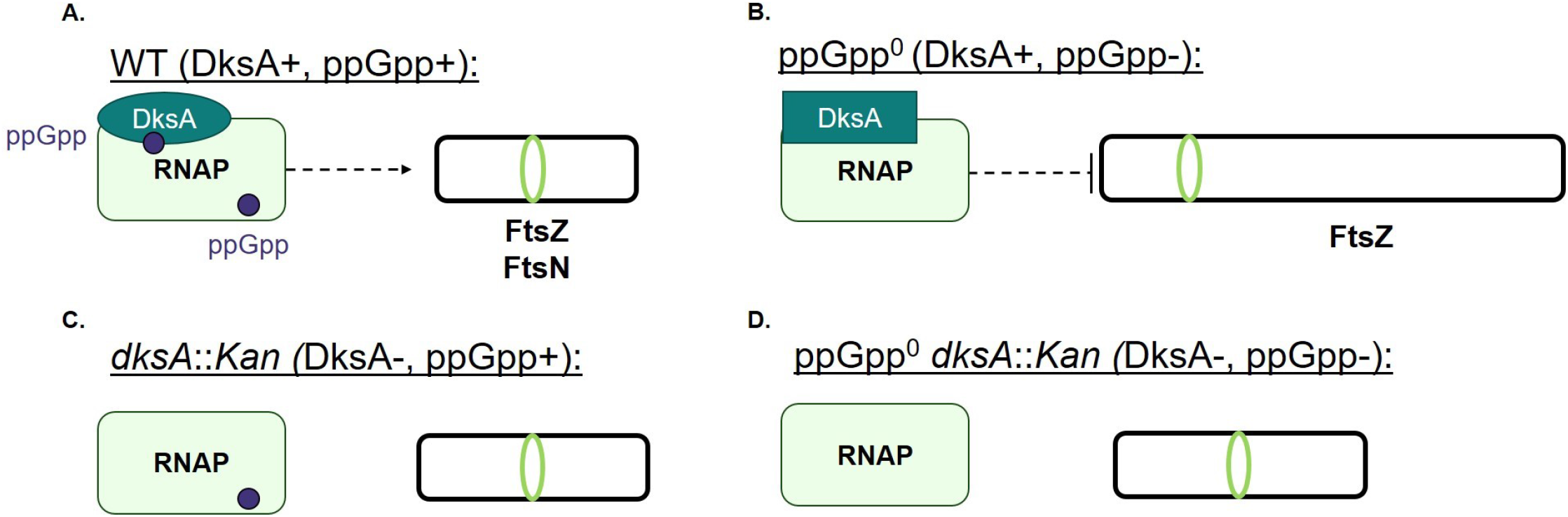
Model of division regulation by ppGpp and DksA. **A.** In wild-type cells, ppGpp and DksA are both available to bind RNAP. The ppGpp/DksA/RNAP complex indirectly activates FtsZ and FtsN, resulting in activation of division and maintenance of wild-type cell size. **B.** In the absence of ppGpp, DksA switches to behaving as an indirect inhibitor of FtsZ, resulting in cell lengthening and filamentation. **C.** In the absence of *dksA*, ppGpp is no longer able to activate division, resulting in lengthening of *dksA*::*Kan* cells compared to wild-type. **D.** In the absence of both ppGpp and DksA, no indirect transcriptional regulation of division occurs. This results in decreased length compared to ppGpp^0^ and similar length to *dksA*::*Kan*.

Our results extend previous work suggesting a link between ppGpp and division. Specifically, we found that ppGpp exerts a positive impact on assembly of the division machinery in a manner that requires interactions with both DksA and RNAP (**Figs. 1, 2, 6**), indicating that ppGpp indirectly regulates division through its effects on transcription. We also established the importance of basal ppGpp levels in maintaining wild-type cell size and modulating division. While previous work showed that cells lacking ppGpp sometimes filament (18, 25), this study established that filaments comprise approximately 20% of ppGpp^0^ populations grown in rich media (**Fig. 7**) (a value that may be slightly underestimated due to the higher probability of long filaments being excluded from cell length calculations due to extending past the field of view or being tangled with other cells). We further uncovered that non-filamentous ppGpp^0^ cells are still longer than wild-type, consistent with a division defect in these cells.

Although collectively our data indicate ppGpp’s effects on division are mediated through transcription (**Fig. 1, 2, 6**), an RNAP_1-2-_ mutant unable to bind ppGpp does not recapitulate the phenotype of a ppGpp^0^ mutant with regard to cell length or hypersensitivity of *ftsZ84* (**Fig. S8**). The RNAP_1-2-_ mutant has a high number of point mutations in multiple polymerase genes: a three codon deletion in *rpoZ* and five different codon substitutions in *rpoC* (all from charged or polar amino acids to alanine) (8, 11). It is possible that these mutations result in pleiotropic changes that mask the effects of loss of ppGpp binding on division. As this is not the only example of phenotypic differences between ppGpp^0^ and RNAP_1-2-_ strains (7, 8), discrepancies between these mutants should be explored further.

Our finding that DksA exhibits opposing effects on division depending on the presence of ppGpp was also unexpected, although there is some precedent for DksA and ppGpp exerting divergent functions in the cell. While ppGpp and DksA are typically thought to work cooperatively to influence transcription, loss of ppGpp or *dksA* leads to opposing transcriptional changes for a subset of genes (15, 48). DksA regulates polyphosphate accumulation in a ppGpp-independent manner (17). ppGpp^0^ and *dksA* mutants have also been reported to exhibit some divergent phenotypes; for example, loss of ppGpp decreases fimbriation and fimbriae-dependent adhesion, whereas loss of *dksA* increases these phenotypes (16, 18). However, to our knowledge, this report is the first example of DksA itself having opposite effects on a single phenotype depending on ppGpp.

These contrasting effects may reflect conformational changes in the RNAP-DksA complex caused by ppGpp binding. Binding of DksA to RNAP results in conformational changes in both, leading to mechanical stress (14). Binding of ppGpp to Site 2 induces further conformational changes that relieve this stress and position the coiled-coil domain of DksA closer to the RNAP active site (14). Together with previous work indicating that the ppGpp and DksA regulons do not completely overlap, the present study suggests that these two conformations of DksA likely have different transcriptional outputs, which lead to opposing effects on cell division.

Our work suggests a mechanism in which the ratio of DksA bound to ppGpp versus DksA unbound from ppGpp modulates cell division and dictates cell length. ppGpp levels vary with the nutritional state, growth phase, and presence of stressors (3), while DksA levels remain constant (and in excess of RNAP levels) throughout growth (12, 65, 66). In the absence of nutritional stress, we propose that steady basal levels of ppGpp allow for the ratio of RNAP bound to DksA/ppGpp vs. DksA alone to be maintained at a certain level, leading to maintenance of cell size. As ppGpp levels increase during stress conditions, more DksA should be in its ppGpp-bound, division activating state, leading to decreased cell length. This effect is likely enhanced by the fact that ppGpp increases DksA’s affinity for RNAP (14, 67). In cells with lower than normal ppGpp levels, or cells lacking ppGpp altogether, the high level of DksA bound to RNAP alone should lead to division inhibition and increased length. Consistent with the observation that length is strongly negatively correlated with ppGpp concentration (42), our model allows for sensitive calibration of cell division in response to changes in ppGpp levels. Why it is advantageous for ppGpp to promote division remains unclear. With limited resources available to devote to cell wall synthesis, it may benefit *E. coli* to prioritize cell division over elongation. This could increase the number of daughter cells generated, strengthening the chances for some members of the population to survive until conditions improve.

Altogether our data indicate that DksA and ppGpp regulate divisome assembly through indirect modulation of FtsZ and FtsN (**Fig. 9**). As the first and last proteins recruited to the divisome, FtsZ and FtsN represent attractive targets for the regulation of division. While overexpression of *ftsZ* reduced the length of both ppGpp^0^ and *dksA*::*Kan* mutants, *ftsN* overexpression solely impacted *dksA*::*Kan* cell size (**Figure 9**). At the same time, the precise mechanism by which ppGpp influences FtsZ and FtsN is unclear. As previously stated, neither DksA nor ppGpp directly transcriptionally regulate *ftsZ* or *ftsN*, nor do they impact FtsZ levels (11, 15, 44, 48, 49) (**Fig. S13**). There are many well-characterized regulators of FtsZ, which control both the placement of the divisome and regulate division in response to stress (28-30, 68-71). FtsN is inhibited in response to DNA damage in *Caulobacter crescentus* (72), and its activity is sensitive to environmental pH in *E. coli* (32). Whether DksA and/or ppGpp affect these known regulatory pathways, or whether they modulate FtsZ and FtsN through a currently unidentified mechanism(s), remains to be determined.

In addition to the open question of precisely how DksA and ppGpp modulate FtsZ and FtsN, this work also raises several additional points for future inquiry. First and foremost, it is unclear whether DksA’s oppositional, ppGpp-dependent effects are limited to regulation of division, or if DksA also exerts divergent regulation on other physiological processes. The prevailing view of DksA as a transcription factor that facilitates regulation by ppGpp may have led phenotypes exerted by DksA in ppGpp’s absence to be overlooked. While our work indicates that the increased average length of ppGpp^0^ cells is due to division inhibition by DksA, it is also unclear why some ppGpp^0^ cells lengthen only slightly, while others form dramatic filaments. Future work focusing on single cells could determine why some ppGpp^0^ cells have slight division defects, while others seem to display a catastrophic failure to divide. Finally, although this work clearly demonstrates that ppGpp regulates division through DksA, the observation that *prelA* still caused a small (albeit non-significant) decrease in length in the *dksA*::*Kan* mutant leaves open the possibility that ppGpp may make additional minor contributions to cell size via other mechanism(s) (**Fig. 1**).

## Materials and Methods

### Bacterial strains, plasmids, and growth conditions

Bacterial strains and plasmids used in this study are detailed in **Table S1**. All experiments were performed in the MG1655 background, referred to as “wild-type.” Alleles of interest were moved between strains via P1 transduction; transductants were confirmed via PCR. *ftsZ84* RNAP_1-2-_ and *ftsZ84* RNAP_1+2+_ strains were generated using a *cat*-linked *ftsZ84* strain. A *cat* insertion in the *ftsZ*-linked *leuO* gene was generated by recombineering using plasmids pKD3 and pKD46 and primers listed in **Table S1** (73). Plasmids *pftsZ* and *pftsQA* were generated from pBS58 (*pftsQAZ*) via Q5 mutagenesis (New England Biolabs, Ipswich, MA) using primers listed in **Table S1**. Primers were acquired from Integrated DNA Technologies (Coralville, IA).

Unless otherwise indicated, all chemicals, media components, and antibiotics were purchased from Sigma-Aldrich (St. Louis, MO). Experiments were performed in LB broth (1% tryptone, 1% NaCl, 0.5% yeast extract) or LBNS broth (1% tryptone, 0.5% yeast extract), as indicated. When indicated, cultures were supplemented with 0.2% glucose or IPTG (see “Fluorescence imaging of division proteins” for concentrations). When selection was necessary, cultures were supplemented with 50 µg/ml kanamycin (Kan), 30 µg/ml chloramphenicol (Cm), 12.5 µg/ml tetracycline (Tet), 100 µg/ml ampicillin (Amp), or 100 µg/ml spectinomycin (Spec).

### Image acquisition

Phase contrast and fluorescence imaging was performed using samples on 1% agarose/PBS pads with an Olympus BX51 microscope equipped with a 100X Plan N (N.A. = 1.25) Ph3 objective (Olympus), X-Cite 120 LED light source (Lumen Dynamics), and an OrcaERG CCD camera (Hammamatsu Photonics) or a Nikon TiE inverted microscope equipped with a 100X Plan N (N.A. = 1.25) objective (Nikon), SOLA SE Light Engine (Lumencor), heated control chamber (OKO Labs), and ORCA-Flash4.0 sCMOS camera (Hammamatsu Photonics). Filter sets for fluorescence were bought from Chroma Technology Corporation. Nikon Elements software (Nikon Instruments) was used for image capture.

### Cell size analysis

Cells were cultured from a single colony and grown to exponential phase (OD_600_ ≥ ∼0.1). Cultures were then back-diluted into fresh media to an OD_600_ of 0.005 and grown to exponential phase (OD_600_ = 0.1-0.2). Cultures were monitored during growth to determine growth rate (see “Growth rate determination”). Cultures were fixed by adding 500 µl culture to 20 µl of 1 M sodium phosphate (pH 7.4) and 100 µl of fixative (16% paraformaldehyde and 8% glutaraldehyde). Samples were incubated at room temperature for 15 min and ice for 30 min. Samples were then stored at 4°C for up to overnight before being pelleted, washed three times in 1 ml PBS, and resuspended in glucose-tris-EDTA (GTE). Samples were stored at 4°C and used for imaging within one week of fixation. Cell length and width were measured from phase contrast images using the FIJI plugin MicrobeJ (52).

### Heat sensitivity experiments

Strains were grown in LB or LB-glucose (supplemented with 100 µg/ml Amp as appropriate) at 30°C until mid-log phase (OD_600_ = ∼0.2-0.6). Cells were pelleted, washed once in LBNS, and resuspended in LBNS to an OD_600_ of 1.0. Cells were serially diluted from 10^-1^ through 10^-6^ in LBNS and 5 µl of each dilution were spot plated on LB (control) and LBNS plates. Plates were incubated at 30°C (LB, LBNS), 37°C, and 42°C (LBNS only) for 20 h prior to imaging.

### Fluorescence imaging of division proteins

Strains producing GFP fusion proteins were grown, sampled, and fixed as in “Cell size analysis.” Strains were grown in the following concentrations of IPTG: *gfp-ftsZ*—1 mM; *gfp-ftsA* and *gfp-ftsL*—100 µM; *gfp-ftsI*—2.5 µM; and *gfp-ftsN*—5 µM, as done previously (32). Fixed samples were imaged within 48 h of fixation. Phase contrast and fluorescence images were acquired on a Nikon TiE inverted microscope. Cell lengths were measured as for “Cell size analysis.” GFP rings were manually counted in FIJI. L/R was calculated using the formula: 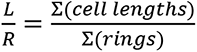.

### Growth rate determination

Growth rates were determined from growth curves of cultures used for fixation (see “Cell size analysis”). OD_600_ was measured every ∼30 min and values were plotted. OD_600_ values within early exponential phase (up to an OD_600_ of 0.2-0.3) were used to calculate growth rate using the Doubling Time Cell Calculator++ (https://doubling-time.com/compute_more.php).

### Western blotting

Strains were grown in LB glucose (wild-type, ppGpp^0^, *dksA*::*Kan*, ppGpp^0^ *dksA*::*Kan*), LB Amp (*prelA’* and *prelA*), or LB Spec (*pftsQAZ*) to mid-log phase (OD_600_ ∼0.3-0.5). Samples were back-diluted to an OD_600_ of 0.01 into the same type of media and grown to an OD_600_ of ∼0.4. Samples were pelleted and stored at -80°C. Samples were thawed on ice and normalized to the same OD_600_ before being resuspended in 4x Laemmli buffer (Bio-Rad, Hercules, CA) with 2-mercaptoethanol and boiled for 10 min. Equivalent volumes of each sample were electrophoresed on a 10% Mini-Protean Precast gel (Bio-Rad) in 25 mM Tris base, 192 mM glycine, 0.1% SDS at 200 V. Proteins were transferred to a PVDF membrane using a Trans-Blot Turbo system (Bio-Rad) using 25 mM Tris base, 192 mM glycine, 20% methanol. Following transfer, membranes were washed for 10 min in PBS. Membranes were agitated in Ponceau stain for 5 min to stain total protein, followed by two washes in PBS 5% acetic acid. Total protein was imaged using an Epson Perfection V600 Photo scanner. Membranes were destained with 0.1 M sodium hydroxide, washed for 5 min in PBS, and blocked for 1 h in 5% milk in PBS. Membranes were incubated overnight in 1:5,000 rabbit α-FtsZ (Cocalico Biologicals, Inc., Stevens, PA) in PBS. Membranes were washed three times for five min in PBS 0.05% Tween, followed by a 1 h incubation in 1:5,000 goat α-rabbit HRP conjugated antibody (Thermo Fisher Scientific, Waltham, MA) in PBS. Membranes were washed three more times in PBS Tween, rinsed twice in PBS, and imaged using Clarity Western ECL Substrate (Bio-Rad) on a LiCor Odyssey imager. Quantitation was determined in FIJI and normalized to Ponceau staining as a total protein loading control (74).

### ppGpp^0^ suppressor tests

ppGpp^0^ strains readily accumulate suppressor mutations in RNAP genes (75). For all experiments using ppGpp^0^ and its derivatives, cultures were tested for the presence of suppressors at the same time that experimental samples were acquired. 500 µl of culture was pelleted and cells were washed once with 1 ml AB media (10 mM ammonium sulfate, 40 mM disodium phosphate, 20 mM monopotassium phosphate, 50 mM sodium chloride, 100 µM calcium chloride, 1 mM magnesium chloride, 3 µM ferric chloride). Cells were resuspended in AB media to an OD_600_ ∼1 and serially diluted from 10^-1^ through 10^-6^. Five µl of each dilution was spot plated on LB and AB 0.2% glucose. Plates were incubated at 37°C overnight (LB) or for up to 48 h (AB-glucose). As ppGpp^0^ is unable to grow on minimal media without amino acids, the frequency of suppressors was determined by dividing the CFU on AB-glucose by the CFU on LB. Data was only included from experiments where the frequency of suppressors was < 10%.

### Quantification and statistical analyses

All experiments were performed with at least three biological replicates. Microscopy experiments were performed using at least 200 cells per replicate, unless otherwise indicated in figure legends. Statistical tests were performed as indicated in figure legends using GraphPad Prism 7.

## Acknowledgements

We would like to thank Dr. Richard Gourse and Wilma Ross for their generous gift of strains and plasmids. We would like to thank Dr. Jade Wang, Dr. Aude Trinquier, and Leah McKinney, as well as members of the Levin lab, for their feedback and discussion. This work was supported by R35-GM127331 to P.A.L. and F32-GM143886 to S.E.A.

---

**Supplemental Figure S1.**
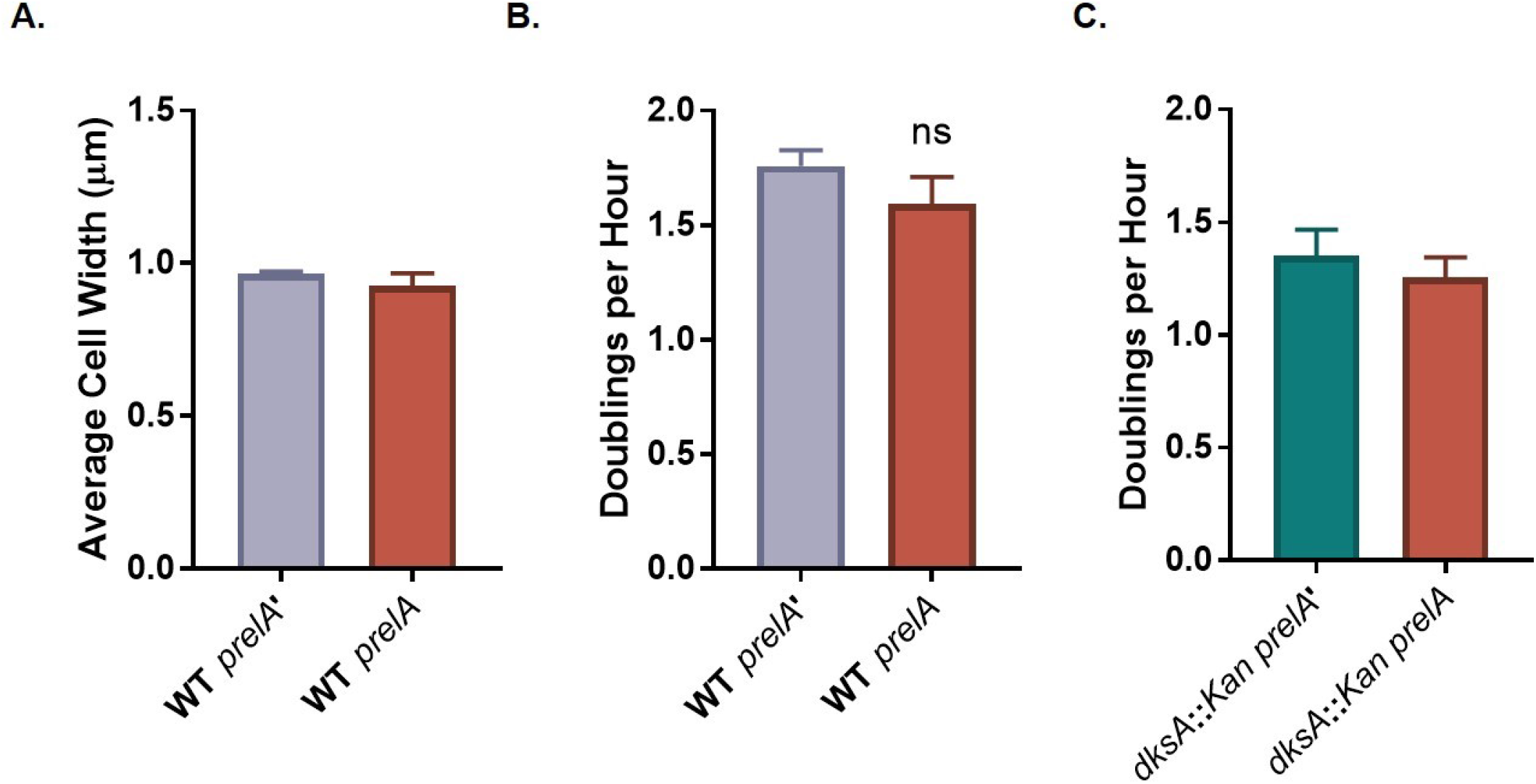
Cell widths and growth rates of strains expressing different levels of ppGpp. **A.** Average width of cells expressing *prelA*. Data represent averages and SDs of at least three independent replicates. Difference is not significant by two-tailed t-test. **B-C.** Growth rates of cells expressing *prelA* in the wild-type (**B**) or *dksA*::*Kan* (**C**) background. Data represent averages and SDs of at least three independent replicates; differences are not significant by two-tailed t-test.

**Supplemental Figure S2.**
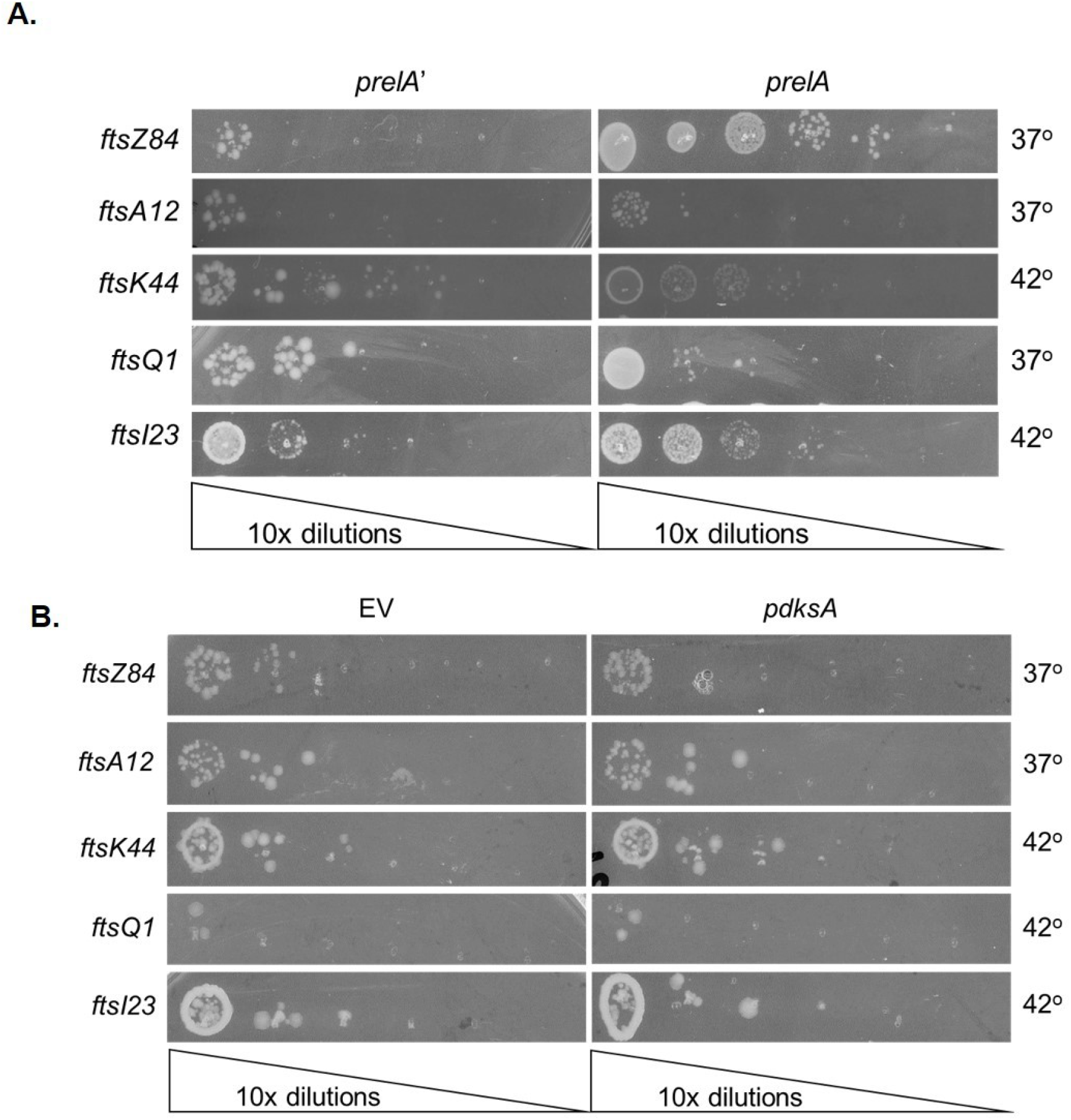
Effect of *relA* and *dksA* overexpression on conditional division mutants. Effect of overexpression of *relA* (**A**) or *dksA* (**B**) on growth of heat-sensitive division mutants at restrictive temperatures. Data shown are representative images of three biological replicates.

**Supplemental Figure S3.**
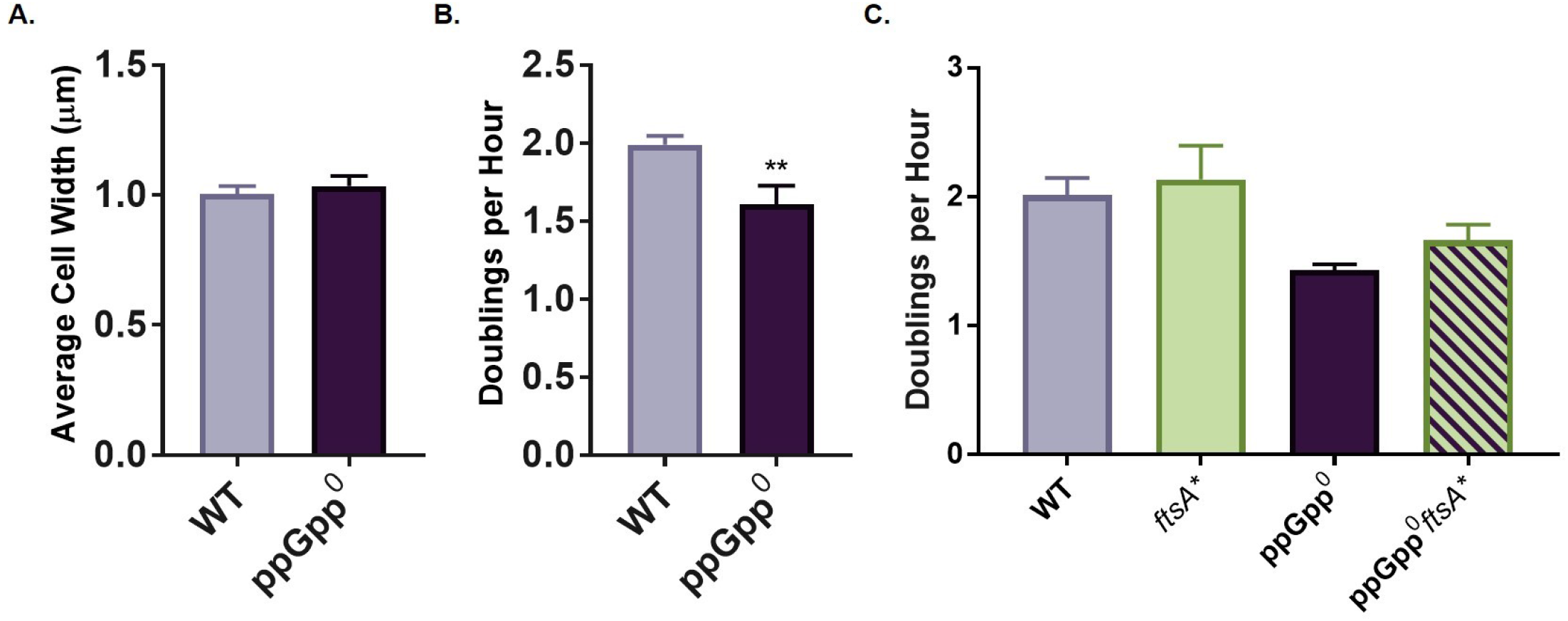
Cell widths and growth rates of ppGpp^0^. **A.** Average widths of cells lacking ppGpp. Data represent averages and SDs of at least three independent replicates. Differences are not significant by two-tailed t-test. **B.** Growth rates of cells lacking ppGpp. Data represent averages and SDs of at least three independent replicates (**, *P* ≤ 0.01 by two-tailed t-test). **C.** Growth rates of ppGpp^0^ and ppGpp^0^ *ftsA\** strains. Data represent averages and SDs of at least three independent replicates. Differences are not significant by one-way ANOVA with Tukey’s multiple comparison test.

**Supplemental Figure S4.**
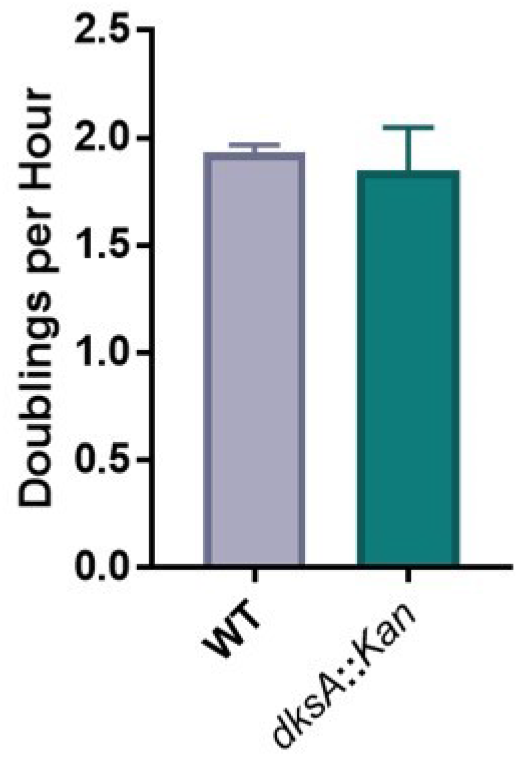
Growth rate of *dksA* mutant. Data represent averages and SDs of three independent replicates. Differences are not significantly different by two-tailed t-test.

**Supplemental Figure S5.**
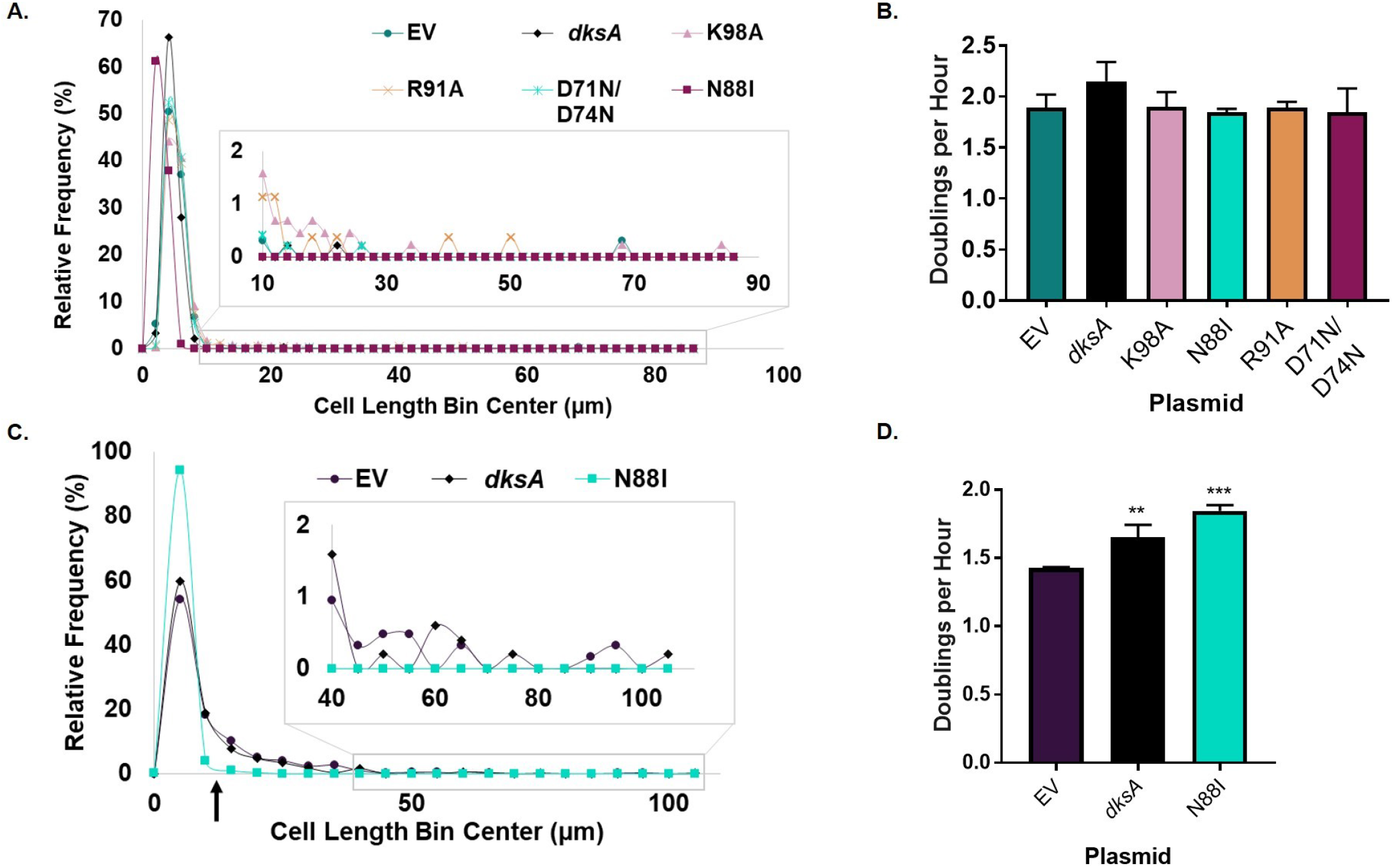
Growth rates and single cell sizes for *dksA*::*Kan* and ppGpp^0^ strains expressing different *dksA* alleles. **A.** Frequency distribution of individual cell lengths for *dksA*::*Kan* complemented with different *dksA* alleles. N > 300 cells from a single representative experiment (bin width = 2 µm). **B.** Expression of different *dksA* alleles from a plasmid has no effect on growth rates of a *dksA*::*Kan* mutant. Data represent averages and SDs of three biological replicates (differences relative to EV are not significant by one-way ANOVA with Dunnett’s post-test). **C.** Frequency distribution of individual cell sizes show that *dksA*_N88I_ reduces length and largely eliminates filamentation in ppGpp^0^. N > 500 cells from a single representative experiment (bin width = 5 µm). The black arrow indicates the approximate location of the cut-off for filamentous cells (12.4 µm, 3x wild-type cell length in Fig. 3C). **D.** Expression of *dksA* and *dksA*_N88I_ increases the growth rate of ppGpp^0^ cells. Data represent averages and SDs of three biological replicates (**, *P* ≤ 0.01; ***, *P* ≤ 0.001 relative to EV by one-way ANOVA with Dunnett’s post-test).

**Supplemental Figure S6.**
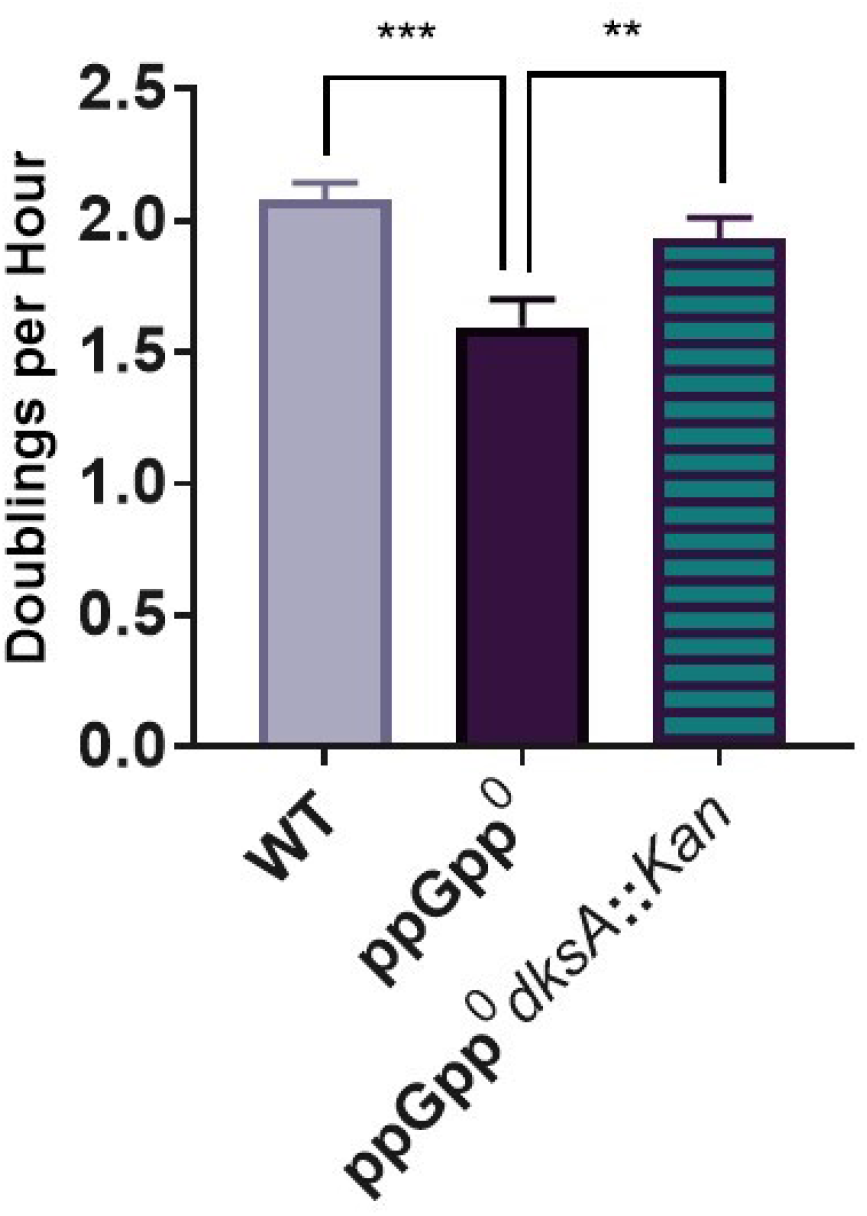
Deletion of *dksA* increases growth rate of ppGpp^0^. Data represent averages and SDs of three biological replicates (**, *P* ≤ 0.01; ***, *P* ≤ 0.001 relative to ppGpp^0^ by one-way ANOVA with Dunnett’s post-test).

**Supplemental Figure S7.**
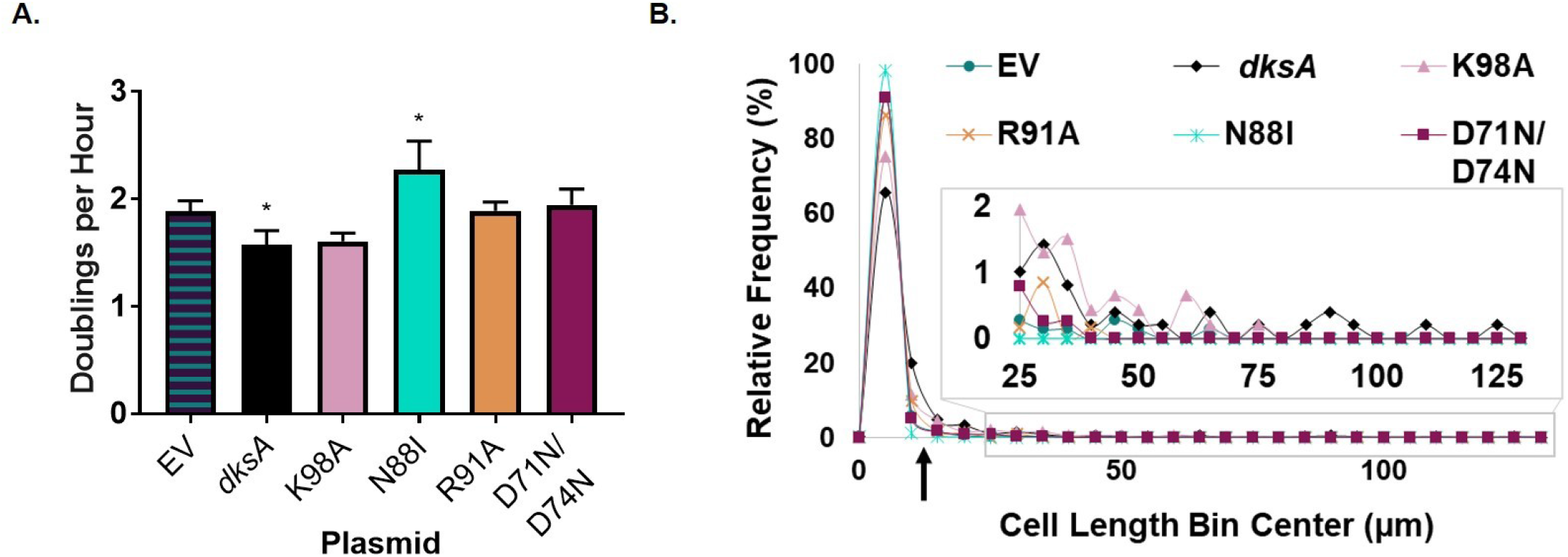
Growth rates and individual cell sizes of ppGpp^0^ *dksA*::*Kan* expressing different *dksA* alleles. **A.** Growth rates of ppGpp^0^ *dksA*::*Kan* mutants complemented with different *dksA* alleles. Complementation with wild-type *dksA* reduces the growth rate, while complementation with *dksA*_N88I_ leads to increased growth rate. Data represent averages and SDs of three independent replicates (*, *P* ≤ 0.05 relative to EV by one-way ANOVA with Dunnett’s post-test). **B.** Frequency distribution of individual cell lengths for ppGpp^0^ *dksA*::*Kan* mutants complemented with different *dksA* alleles. N > 300 cells from a single representative experiment (bin width = 5 µm). The arrow indicates the approximate location of the cut-off for filamentous cells (12.2 µm, 3x wild-type length from Fig. 7A).

**Supplemental Figure S8.**
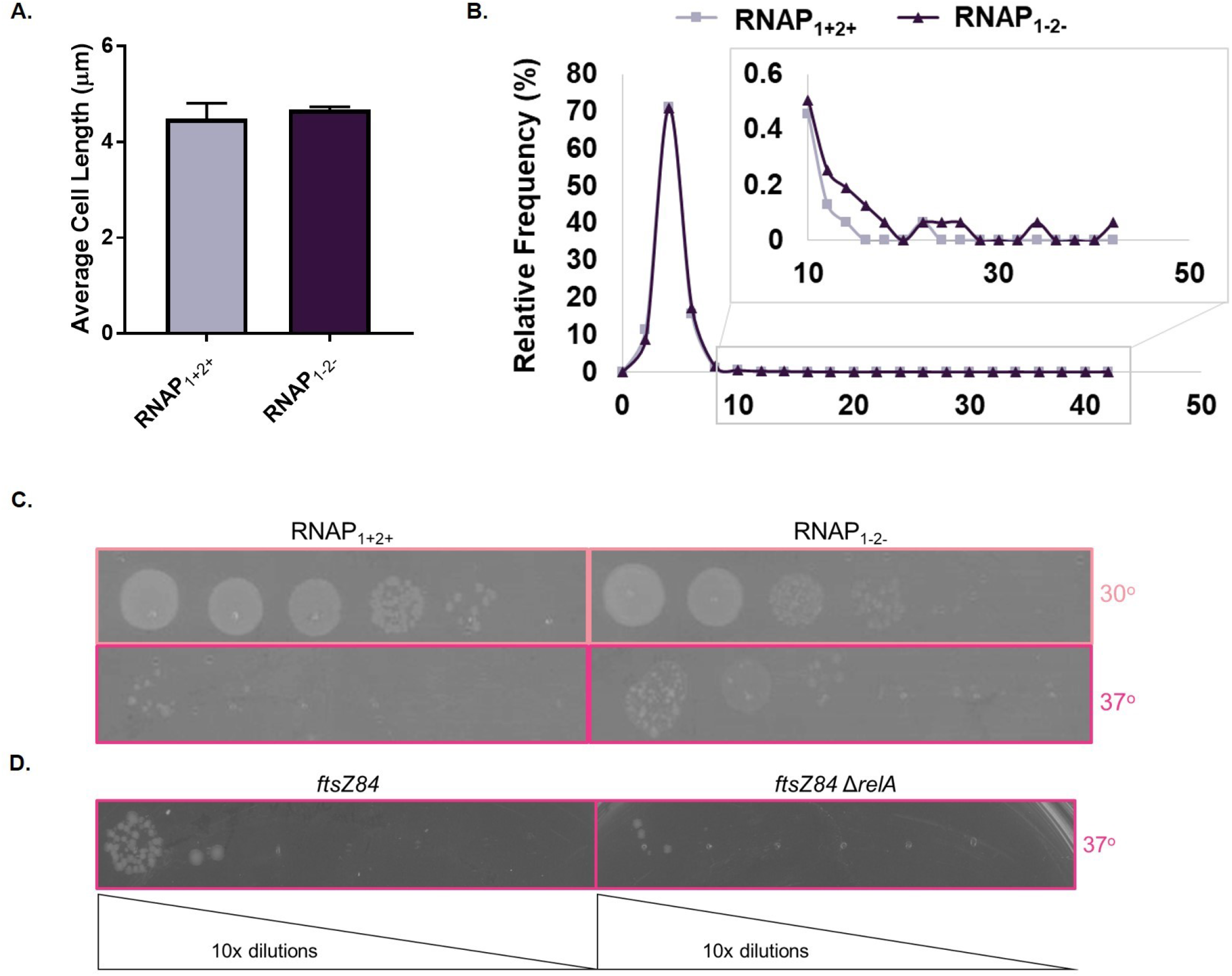
Mutations blocking ppGpp binding to RNAP do not affect cell length. **A.** There is no significant difference in cell length between an RNAP_1-2-_ mutant and an RNAP_1+2+_ control by two-tailed t-test. Data represent averages and SDs of three independent replicates. **B.** Distribution of individual cell lengths reveal that RNAP_1-2-_ cells have very similar lengths to RNAP_1+2+_ cells. N > 1500 cells from a single representative experiment (bin width = 2 µm). **C.** An RNAP_1-2-_ *ftsZ84* exhibits a slight decrease in CFUs at 30°C and an increase in CFUs at 37°C compared to an RNAP_1+2+_ *ftsZ84* control. A representative image from three biological replicates is shown. **D.** Deletion of *relA* does not affect growth of *ftsZ84* at the non-permissive temperature of 37°C. A representative image from three biological replicates is shown.

**Supplemental Figure S9.**
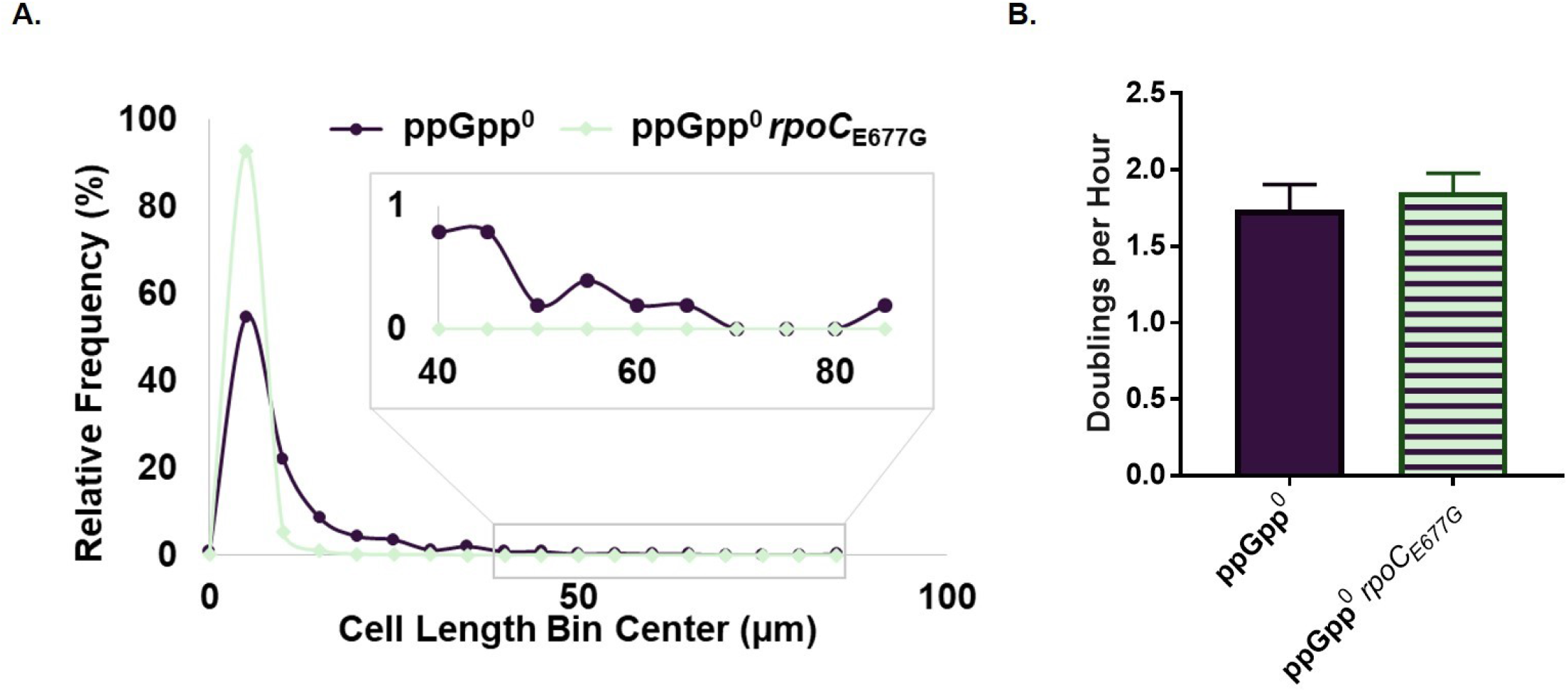
Growth rates and individual cell sizes of ppGpp^0^ expressing different *dksA* or *rpoC* alleles. **A.** Frequency distribution of individual cell lengths reveal that the *rpoC*_E677G_ mutation reduces length and filamentation of ppGpp^0^ cells. N > 500 cells from a representative biological replicate (bin width = 5 µm). **B.** The *rpoC*_E677G_ mutation does not affect growth rate of ppGpp^0^ cells. Data represent averages and SDs of three independent replicates (differences are not significant by two-tailed t-test).

**Supplemental Figure S10.**
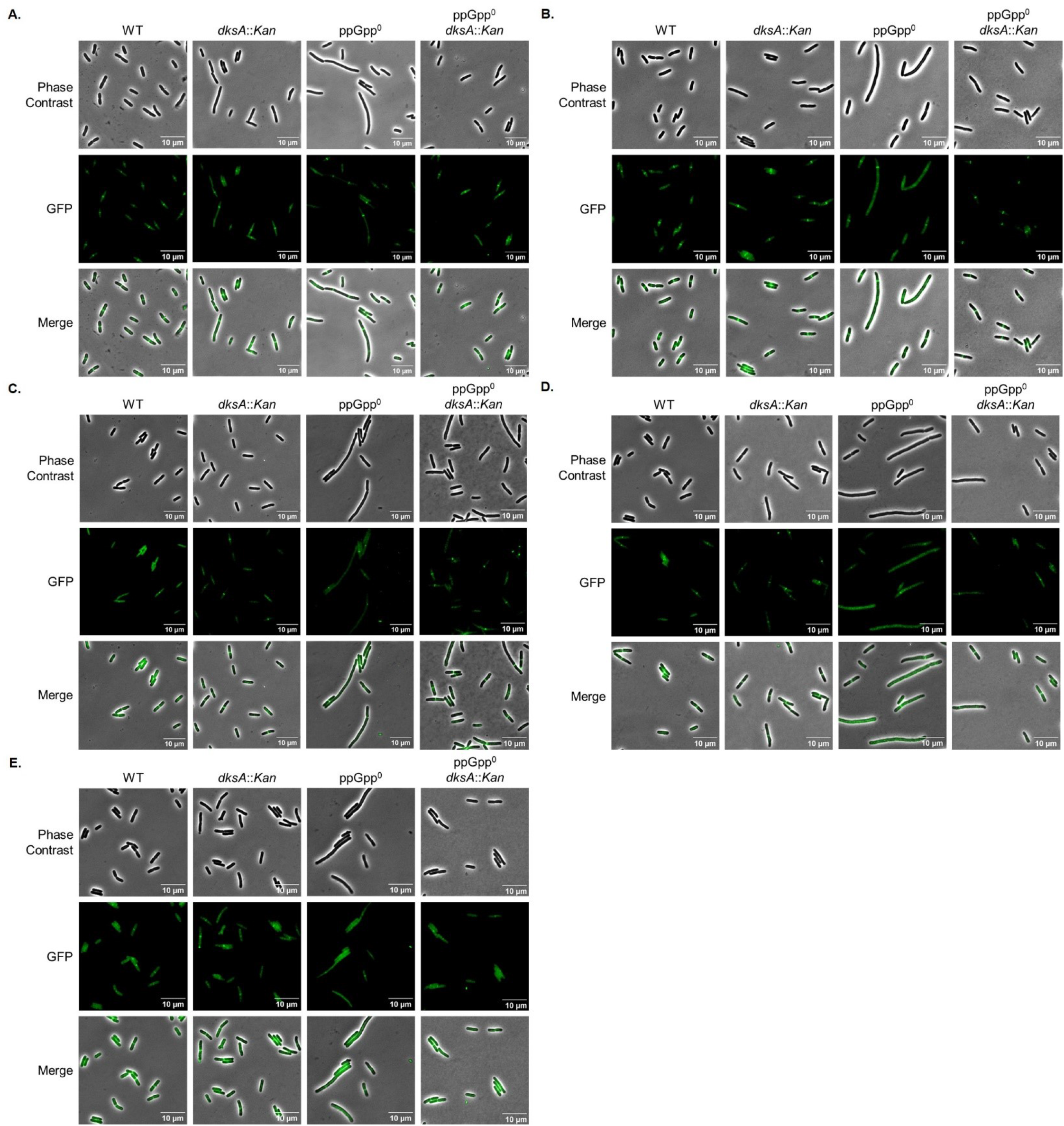
ppGpp^0^ cells exhibit decreased recruitment of division proteins. Phase contrast and fluorescence images of cells expressing GFP-FtsZ (**A**), GFP-FtsA (**B**), GFP-FtsL (**C**), GFP-FtsI (**D**), or GFP-FtsN (**E**) are shown. Images are representative of at least three biological replicates.

**Supplemental Figure S11.**
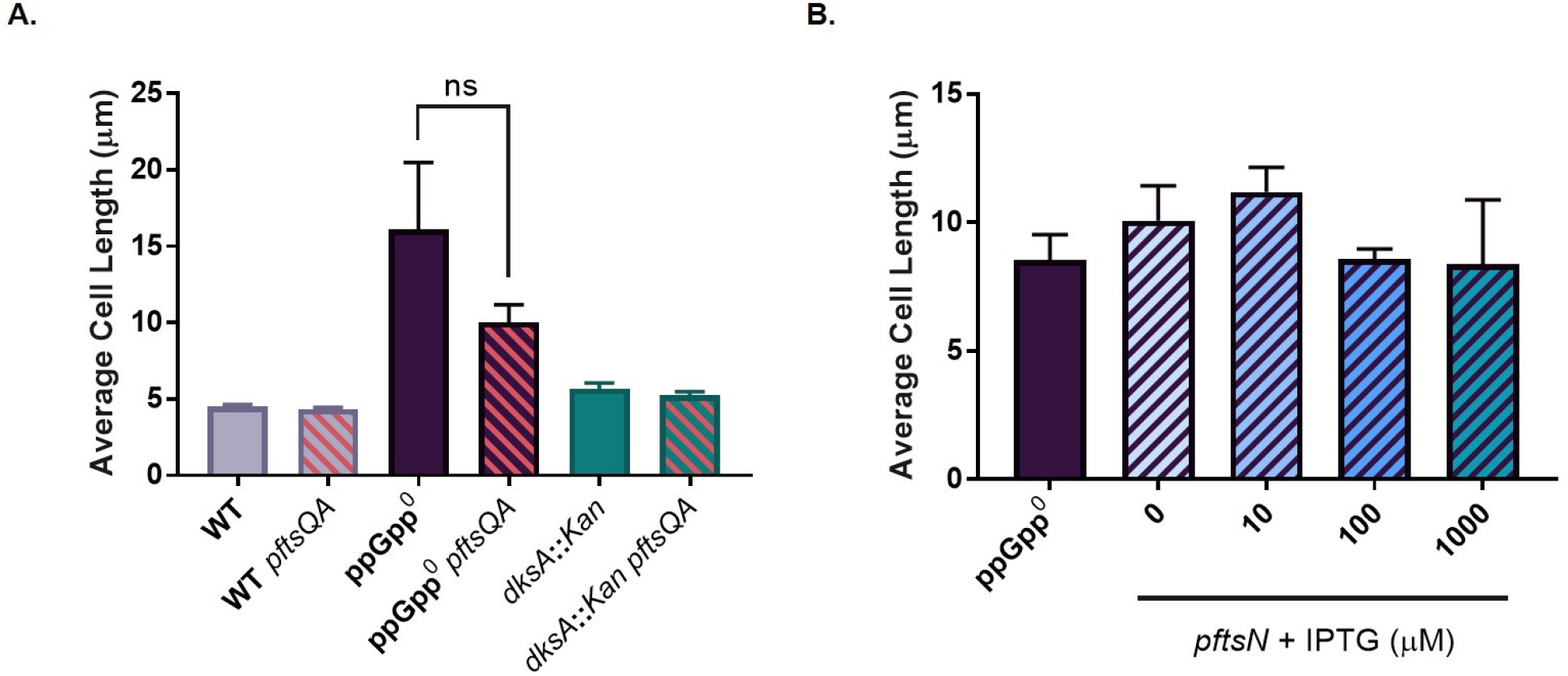
*ftsQA* and *ftsN* do not reduce the length of ppGpp^0^ cells. **A.** Overexpression of *ftsQA* does not lead to differences in cell length in any background. Data shown represent averages and SDs from three biological replicates. Differences are not significant by two-tailed t-test. **B.** Overexpression of *gfp-ftsN* from an IPTG-inducible plasmid has no effect on length of ppGpp^0^ cells. Data shown represent averages and SDs from three biological replicates; differences are not significant by one-way ANOVA with Dunnett’s post-test relative to ppGpp^0^.

**Supplemental Figure S12.**
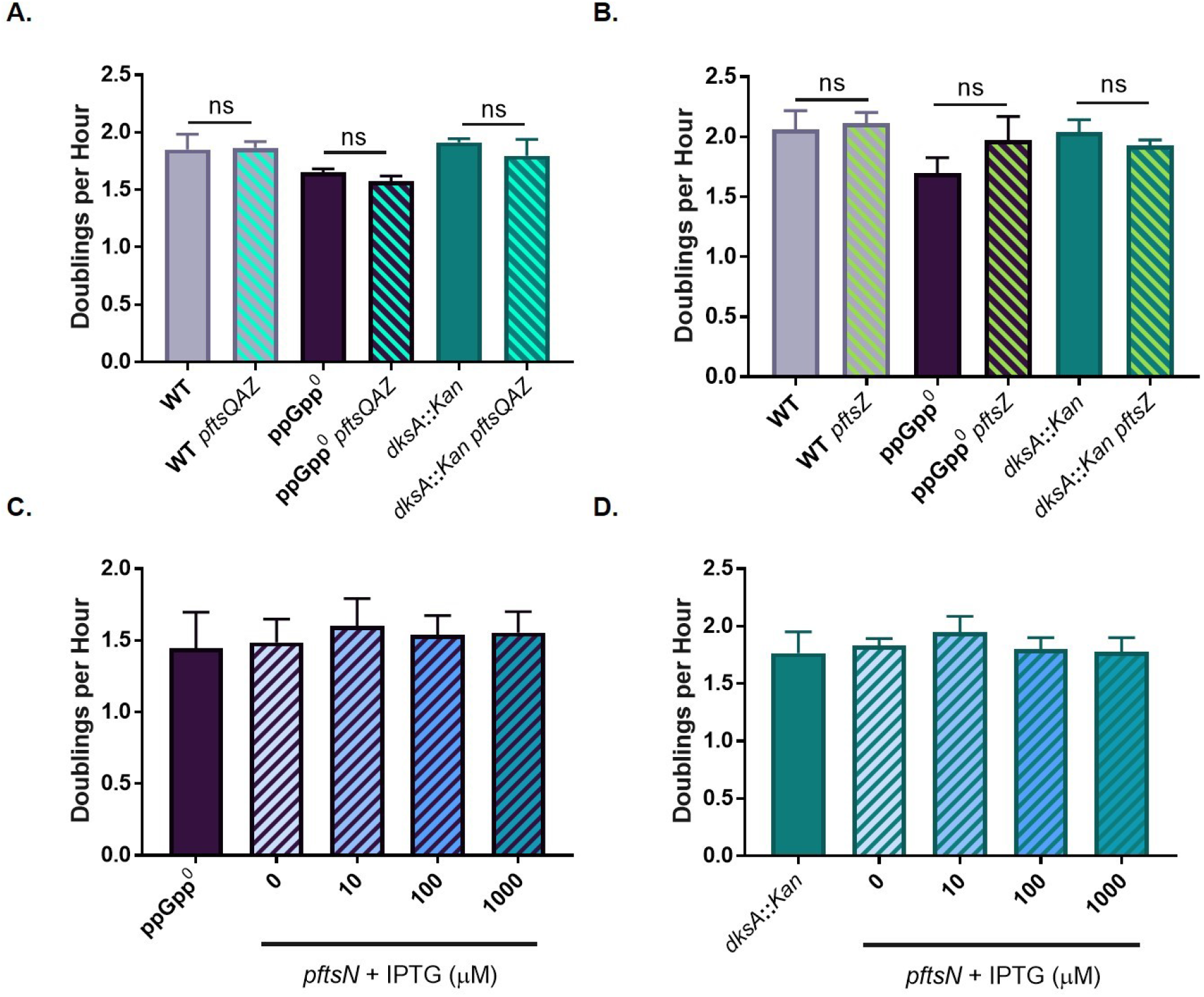
Overexpression of division proteins does not lead to changes in growth rate. **A-B.** No significant differences in growth rates were observed for strains overexpressing *ftsQAZ* (**A**) or *ftsZ* (**B**). Data represent averages and SDs of three biological replicates (ns, not significant by two-tailed t-test). **C-D.** No significant differences in growth rates were observed for ppGpp^0^ (**C**) or *dksA*::*Kan* (**D**) overexpressing *gfp-ftsN*. Data represent averages and SDs of three biological replicates; differences are not significant relative to vector-free controls by one-way ANOVA with Dunnett’s post-test.

**Supplemental Figure S13.**
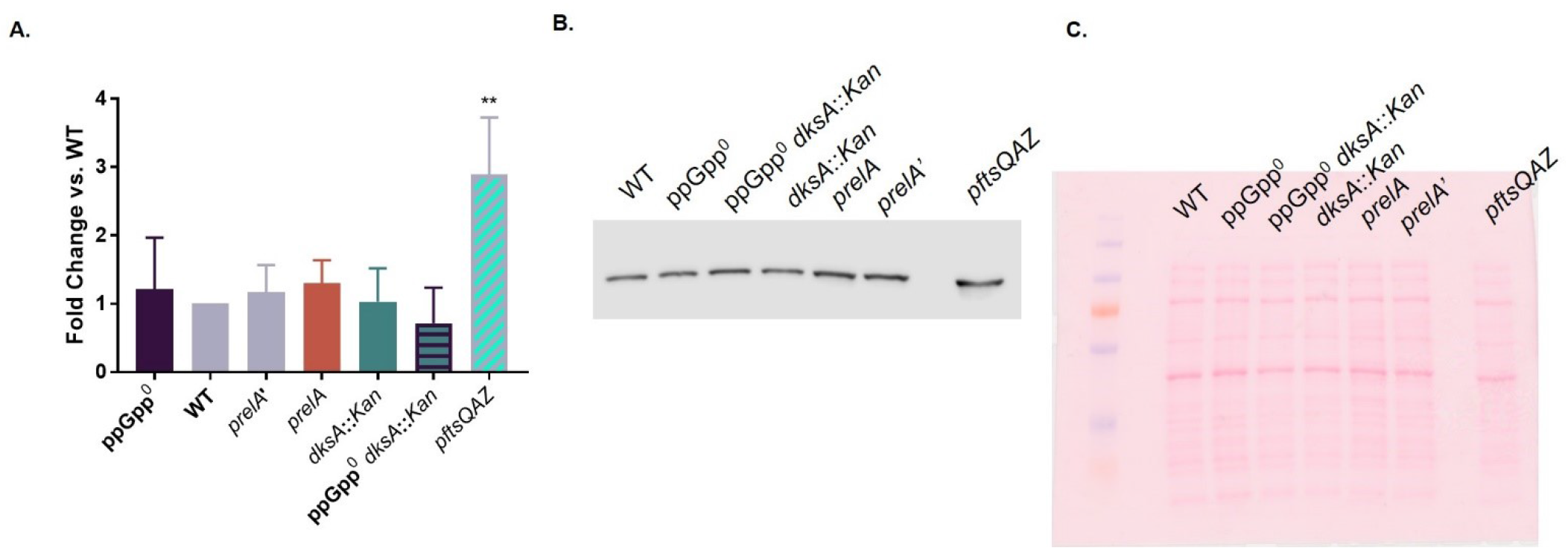
DksA and ppGpp do not affect FtsZ levels. **A.** Strains with varying levels of *dksA* or ppGpp have no change in FtsZ levels compared to wild-type. FtsZ concentrations were normalized to total protein and to wild-type. Data represent averages and SDs of three independent replicates (**, *P* ≤ 0.01 relative to wild-type by one-way ANOVA with Dunnett’s post-test). **B.** A representative FtsZ immunoblot is shown. **C.** A representative Ponceau stain for total protein is shown.

## References

1. Ronneau S, Hallez R. 2019. Make and break the alarmone: regulation of (p)ppGpp synthetase/hydrolase enzymes in bacteria. FEMS Microbiol Rev 43:389–400.

2. Kudrin P, Dzhygyr I, Ishiguro K, Beljantseva J, Maksimova E, Oliveira SRA, Varik V, Payoe R, Konevega AL, Tenson T, Suzuki T, Hauryliuk V. 2018. The ribosomal A-site finger is crucial for binding and activation of the stringent factor RelA. Nucleic Acids Res 46:1973–1983.

3. Steinchen W, Zegarra V, Bange G. 2020. (p)ppGpp: Magic Modulators of Bacterial Physiology and Metabolism. Front Microbiol 11:2072.

4. Germain E, Guiraud P, Byrne D, Douzi B, Djendli M, Maisonneuve E. 2019. YtfK activates the stringent response by triggering the alarmone synthetase SpoT in Escherichia coli. Nature Communications 10:5763.

5. Dalebroux ZD, Swanson MS. 2012. ppGpp: magic beyond RNA polymerase. Nat Rev Microbiol 10:203–12.

6. Lee JW, Park YH, Seok YJ. 2018. Rsd balances (p)ppGpp level by stimulating the hydrolase activity of SpoT during carbon source downshift in Escherichia coli. Proc Natl Acad Sci U S A 115:E6845–e6854.

7. Fernández-Coll L, Cashel M. 2020. Possible Roles for Basal Levels of (p)ppGpp: Growth Efficiency Vs. Surviving Stress. Front Microbiol 11:592718.

8. Ross W, Sanchez-Vazquez P, Chen AY, Lee JH, Burgos HL, Gourse RL. 2016. ppGpp Binding to a Site at the RNAP-DksA Interface Accounts for Its Dramatic Effects on Transcription Initiation during the Stringent Response. Mol Cell 62:811–823.

9. Ross W, Vrentas CE, Sanchez-Vazquez P, Gaal T, Gourse RL. 2013. The magic spot: a ppGpp binding site on E. coli RNA polymerase responsible for regulation of transcription initiation. Mol Cell 50:420–9.

10. Myers AR, Thistle DP, Ross W, Gourse RL. 2020. Guanosine Tetraphosphate Has a Similar Affinity for Each of Its Two Binding Sites on Escherichia coli RNA Polymerase. Front Microbiol 11:587098.

11. Sanchez-Vazquez P, Dewey CN, Kitten N, Ross W, Gourse RL. 2019. Genome-wide effects on *Escherichia coli* transcription from ppGpp binding to its two sites on RNA polymerase. Proceedings of the National Academy of Sciences 116:8310–8319.

12. Paul BJ, Barker MM, Ross W, Schneider DA, Webb C, Foster JW, Gourse RL. 2004. DksA: A Critical Component of the Transcription Initiation Machinery that Potentiates the Regulation of rRNA Promoters by ppGpp and the Initiating NTP. Cell 118:311–322.

13. Paul BJ, Berkmen MB, Gourse RL. 2005. DksA potentiates direct activation of amino acid promoters by ppGpp. Proceedings of the National Academy of Sciences 102:7823–7828.

14. Molodtsov V, Sineva E, Zhang L, Huang X, Cashel M, Ades SE, Murakami KS. 2018. Allosteric Effector ppGpp Potentiates the Inhibition of Transcript Initiation by DksA. Mol Cell 69:828–839.e5.

15. Aberg A, Fernández-Vázquez J, Cabrer-Panes JD, Sánchez A, Balsalobre C. 2009. Similar and divergent effects of ppGpp and DksA deficiencies on transcription in Escherichia coli. J Bacteriol 191:3226–36.

16. Åberg A, Shingler V, Balsalobre C. 2008. Regulation of the fimB promoter: a case of differential regulation by ppGpp and DksA in vivo. Molecular Microbiology 67:1223–1241.

17. Gray MJ. 2019. Inorganic Polyphosphate Accumulation in Escherichia coli Is Regulated by DksA but Not by (p)ppGpp. J Bacteriol 201.

18. Magnusson LU, Gummesson B, Joksimović P, Farewell A, Nyström T. 2007. Identical, Independent, and Opposing Roles of ppGpp and DksA in *Escherichia coli*. Journal of Bacteriology 189:5193-5202.

19. Gopalkrishnan S, Nicoloff H, Ades SE. 2014. Co-ordinated regulation of the extracytoplasmic stress factor, sigmaE, with other Escherichia coli sigma factors by (p)ppGpp and DksA may be achieved by specific regulation of individual holoenzymes. Molecular Microbiology 93:479–493.

20. Huang C, Meng J, Li W, Chen J. 2022. Similar and Divergent Roles of Stringent Regulator (p)ppGpp and DksA on Pleiotropic Phenotype of Yersinia enterocolitica. Microbiol Spectr 10:e0205522.

21. Wang B, Dai P, Ding D, Del Rosario A, Grant RA, Pentelute BL, Laub MT. 2019. Affinity-based capture and identification of protein effectors of the growth regulator ppGpp. Nat Chem Biol 15:141–150.

22. Wang B, Grant RA, Laub MT. 2020. ppGpp Coordinates Nucleotide and Amino-Acid Synthesis in E. coli During Starvation. Mol Cell 80:29–42.e10.

23. Zhang Y, Zborníková E, Rejman D, Gerdes K. 2018. Novel (p)ppGpp Binding and Metabolizing Proteins of Escherichia coli. mBio 9.

24. Fernández-Coll L, Maciag-Dorszynska M, Tailor K, Vadia S, Levin PA, Szalewska-Palasz A, Cashel M. 2020. The Absence of (p)ppGpp Renders Initiation of Escherichia coli Chromosomal DNA Synthesis Independent of Growth Rates. mBio 11.

25. Xiao H, Kalman M, Ikehara K, Zemel S, Glaser G, Cashel M. 1991. Residual guanosine 3’,5’-bispyrophosphate synthetic activity of relA null mutants can be eliminated by spoT null mutations. J Biol Chem 266:5980–90.

26. Nazir A, Harinarayanan R. 2015. Inactivation of Cell Division Protein FtsZ by SulA Makes Lon Indispensable for the Viability of a ppGpp0 Strain of Escherichia coli. J Bacteriol 198:688–700.

27. Levin PA, Janakiraman A. 2021. Localization, Assembly, and Activation of the Escherichia coli Cell Division Machinery. EcoSal Plus 9:eESP00222021.

28. Bi E, Lutkenhaus J. 1993. Cell division inhibitors SulA and MinCD prevent formation of the FtsZ ring. J Bacteriol 175:1118–25.

29. Hill NS, Buske PJ, Shi Y, Levin PA. 2013. A Moonlighting Enzyme Links Escherichia coli Cell Size with Central Metabolism. PLOS Genetics 9:e1003663.

30. Bernhardt TG, de Boer PAJ. 2005. SlmA, a Nucleoid-Associated, FtsZ Binding Protein Required for Blocking Septal Ring Assembly over Chromosomes in E. coli. Molecular Cell 18:555–564.

31. Camberg JL, Hoskins JR, Wickner S. 2009. ClpXP protease degrades the cytoskeletal protein, FtsZ, and modulates FtsZ polymer dynamics. Proceedings of the National Academy of Sciences 106:10614–10619.

32. Mueller EA, Westfall CS, Levin PA. 2020. pH-dependent activation of cytokinesis modulates Escherichia coli cell size. PLoS Genet 16:e1008685.

33. Chen JC, Weiss DS, Ghigo JM, Beckwith J. 1999. Septal localization of FtsQ, an essential cell division protein in Escherichia coli. J Bacteriol 181:521–30.

34. Begg KJ, Dewar SJ, Donachie WD. 1995. A new Escherichia coli cell division gene, ftsK. J Bacteriol 177:6211–22.

35. Arjes HA, Lai B, Emelue E, Steinbach A, Levin PA. 2015. Mutations in the bacterial cell division protein FtsZ highlight the role of GTP binding and longitudinal subunit interactions in assembly and function. BMC Microbiol 15:209.

36. Weiss DS, Chen JC, Ghigo JM, Boyd D, Beckwith J. 1999. Localization of FtsI (PBP3) to the septal ring requires its membrane anchor, the Z ring, FtsA, FtsQ, and FtsL. J Bacteriol 181:508–20.

37. Dai K, Xu Y, Lutkenhaus J. 1993. Cloning and characterization of ftsN, an essential cell division gene in Escherichia coli isolated as a multicopy suppressor of ftsA12(Ts). J Bacteriol 175:3790–7.

38. Geissler B, Elraheb D, Margolin W. 2003. A gain-of-function mutation in ftsA bypasses the requirement for the essential cell division gene zipA in Escherichia coli. Proc Natl Acad Sci U S A 100:4197–202.

39. Geissler B, Margolin W. 2005. Evidence for functional overlap among multiple bacterial cell division proteins: compensating for the loss of FtsK. Molecular Microbiology 58:596–612.

40. Schoenemann KM, Krupka M, Rowlett VW, Distelhorst SL, Hu B, Margolin W. 2018. Gain-of-function variants of FtsA form diverse oligomeric structures on lipids and enhance FtsZ protofilament bundling. Mol Microbiol 109:676–693.

41. Powell BS, Court DL. 1998. Control of ftsZ expression, cell division, and glutamine metabolism in Luria-Bertani medium by the alarmone ppGpp in Escherichia coli. J Bacteriol 180:1053–62.

42. Büke F, Grilli J, Cosentino Lagomarsino M, Bokinsky G, Tans SJ. 2021. ppGpp is a bacterial cell size regulator. Curr Biol doi:10.1016/j.cub.2021.12.033.

43. Schreiber G, Ron EZ, Glaser G. 1995. ppGpp-mediated regulation of DNA replication and cell division in Escherichia coli. Curr Microbiol 30:27–32.

44. Navarro F, Robin A, D’Ari R, Joseleau-Petit D. 1998. Analysis of the effect of ppGpp on the ftsQAZ operon in Escherichia coli. Mol Microbiol 29:815–23.

45. Lai GC, Cho H, Bernhardt TG. 2017. The mecillinam resistome reveals a role for peptidoglycan endopeptidases in stimulating cell wall synthesis in Escherichia coli. PLoS Genet 13:e1006934.

46. Kocaoglu O, Carlson EE. 2015. Profiling of β-Lactam Selectivity for Penicillin-Binding Proteins in Escherichia coli Strain DC2. Antimicrobial Agents and Chemotherapy 59:2785–2790.

47. Kruse T, Bork-Jensen J, Gerdes K. 2005. The morphogenetic MreBCD proteins of Escherichia coli form an essential membrane-bound complex. Mol Microbiol 55:78–89.

48. Vinella D, Potrykus K, Murphy H, Cashel M. 2012. Effects on growth by changes of the balance between GreA, GreB, and DksA suggest mutual competition and functional redundancy in Escherichia coli. J Bacteriol 194:261–73.

49. Traxler MF, Summers SM, Nguyen HT, Zacharia VM, Hightower GA, Smith JT, Conway T. 2008. The global, ppGpp-mediated stringent response to amino acid starvation in Escherichia coli. Mol Microbiol 68:1128–48.

50. Svitil AL, Cashel M, Zyskind JW. 1993. Guanosine tetraphosphate inhibits protein synthesis in vivo. A possible protective mechanism for starvation stress in Escherichia coli. J Biol Chem 268:2307–11.

51. Heath RJ, Jackowski S, Rock CO. 1994. Guanosine tetraphosphate inhibition of fatty acid and phospholipid synthesis in Escherichia coli is relieved by overexpression of glycerol-3-phosphate acyltransferase (plsB). J Biol Chem 269:26584–90.

52. Ducret A, Quardokus EM, Brun YV. 2016. MicrobeJ, a tool for high throughput bacterial cell detection and quantitative analysis. Nature Microbiology 1:16077.

53. Lutkenhaus JF, Wolf-Watz H, Donachie WD. 1980. Organization of genes in the ftsA-envA region of the Escherichia coli genetic map and identification of a new fts locus (ftsZ). J Bacteriol 142:615–20.

54. Herricks JR, Nguyen D, Margolin W. 2014. A thermosensitive defect in the ATP binding pocket of FtsA can be suppressed by allosteric changes in the dimer interface. Molecular microbiology 94:713–727.

55. Begg KJ, Hatfull GF, Donachie WD. 1980. Identification of new genes in a cell envelope-cell division gene cluster of Escherichia coli: cell division gene ftsQ. J Bacteriol 144:435–7.

56. Begg KJ, Donachie WD. 1985. Cell shape and division in Escherichia coli: experiments with shape and division mutants. J Bacteriol 163:615–22.

57. Bernard CS, Sadasivam M, Shiomi D, Margolin W. 2007. An altered FtsA can compensate for the loss of essential cell division protein FtsN in Escherichia coli. Mol Microbiol 64:1289–305.

58. Radler P, Baranova N, Caldas P, Sommer C, López-Pelegrín M, Michalik D, Loose M. 2022. In vitro reconstitution of Escherichia coli divisome activation. Nature Communications 13:2635.

59. Männik J, Pichoff S, Lutkenhaus J, Männik J. 2022. Cell Cycle-Dependent Recruitment of FtsN to the Divisome in Escherichia coli. mBio doi:10.1128/mbio.02017-22:e0201722.

60. Blankschien MD, Lee JH, Grace ED, Lennon CW, Halliday JA, Ross W, Gourse RL, Herman C. 2009. Super DksAs: substitutions in DksA enhancing its effects on transcription initiation. Embo j 28:1720–31.

61. Parshin A, Shiver AL, Lee J, Ozerova M, Schneidman-Duhovny D, Gross CA, Borukhov S. 2015. DksA regulates RNA polymerase in *Escherichia coli*through a network of interactions in the secondary channel that includes Sequence Insertion 1. Proceedings of the National Academy of Sciences 112:E6862-E6871.

62. Lee J-H, Lennon CW, Ross W, Gourse RL. 2012. Role of the Coiled-Coil Tip of Escherichia coli DksA in Promoter Control. Journal of Molecular Biology 416:503–517.

63. Satory D, Halliday JA, Sivaramakrishnan P, Lua RC, Herman C. 2013. Characterization of a novel RNA polymerase mutant that alters DksA activity. J Bacteriol 195:4187–94.

64. Bi E, Lutkenhaus J. 1990. FtsZ regulates frequency of cell division in Escherichia coli. Journal of Bacteriology 172:2765–2768.

65. Chandrangsu P, Lemke JJ, Gourse RL. 2011. The dksA promoter is negatively feedback regulated by DksA and ppGpp. Mol Microbiol 80:1337–48.

66. Rutherford ST, Lemke JJ, Vrentas CE, Gaal T, Ross W, Gourse RL. 2007. Effects of DksA, GreA, and GreB on transcription initiation: insights into the mechanisms of factors that bind in the secondary channel of RNA polymerase. J Mol Biol 366:1243–57.

67. Shin Y, Qayyum MZ, Pupov D, Esyunina D, Kulbachinskiy A, Murakami KS. 2021. Structural basis of ribosomal RNA transcription regulation. Nat Commun 12:528.

68. Buss JA, Peters NT, Xiao J, Bernhardt TG. 2017. ZapA and ZapB form an FtsZ-independent structure at midcell. Mol Microbiol 104:652–663.

69. Marteyn BS, Karimova G, Fenton AK, Gazi AD, West N, Touqui L, Prevost MC, Betton JM, Poyraz O, Ladant D, Gerdes K, Sansonetti PJ, Tang CM. 2014. ZapE is a novel cell division protein interacting with FtsZ and modulating the Z-ring dynamics. mBio 5:e00022–14.

70. Camberg JL, Hoskins JR, Wickner S. 2011. The interplay of ClpXP with the cell division machinery in Escherichia coli. J Bacteriol 193:1911–8.

71. Männik J, Walker BE, Männik J. 2018. Cell cycle-dependent regulation of FtsZ in Escherichia coli in slow growth conditions. Mol Microbiol 110:1030–1044.

72. Modell JW, Kambara TK, Perchuk BS, Laub MT. 2014. A DNA damage-induced, SOS-independent checkpoint regulates cell division in Caulobacter crescentus. PLoS Biol 12:e1001977.

73. Datsenko KA, Wanner BL. 2000. One-step inactivation of chromosomal genes in Escherichia coli K-12 using PCR products. Proceedings of the National Academy of Sciences of the United States of America 97:6640–6645.

74. Schindelin J, Arganda-Carreras I, Frise E, Kaynig V, Longair M, Pietzsch T, Preibisch S, Rueden C, Saalfeld S, Schmid B, Tinevez J-Y, White DJ, Hartenstein V, Eliceiri K, Tomancak P, Cardona A. 2012. Fiji: an open-source platform for biological-image analysis. Nature Methods 9:676-682.

75. Murphy H, Cashel M. 2003. Isolation of RNA Polymerase Suppressors of a (p)ppGpp Deficiency, p 596-601, Methods in Enzymology, vol 371. Academic Press.

76. Fürste JP, Pansegrau W, Frank R, Blöcker H, Scholz P, Bagdasarian M, Lanka E. 1986. Molecular cloning of the plasmid RP4 primase region in a multi-host-range tacP expression vector. Gene 48:119–31.

77. Masui Y, Mizuno T, Inouye M. 1984. Novel High-level Expression Cloning Vehicles: 104-fold Amplification of Escherichia coli Minor Protein. Bio/Technology 2:81–85.

78. Gerding MA, Liu B, Bendezú FO, Hale CA, Bernhardt TG, de Boer PA. 2009. Self-enhanced accumulation of FtsN at Division Sites and Roles for Other Proteins with a SPOR domain (DamX, DedD, and RlpA) in Escherichia coli cell constriction. J Bacteriol 191:7383–401.

79. Hale CA, de Boer PA. 2002. ZipA is required for recruitment of FtsK, FtsQ, FtsL, and FtsN to the septal ring in Escherichia coli. J Bacteriol 184:2552–6.

80. Perederina A, Svetlov V, Vassylyeva MN, Tahirov TH, Yokoyama S, Artsimovitch I, Vassylyev DG. 2004. Regulation through the secondary channel--structural framework for ppGpp-DksA synergism during transcription. Cell 118:297–309.

