## Supplemental Table 1 for "The transcription factor DksA exerts opposing effects on cell division depending on the presence of ppGpp"

**Table S1.** Bacterial strains, plasmids, and primers used in this study.

**Strains**

| Designation | Genotype | Source |
|---|---|---|
| MG1655 | *rph1 ilvG rfb-50 λ- F-* | (1) |
| CSW808 | MG1655 Δ*relA spoT*::*cat* (ppGpp$^0$) | This study |
| JW0141-1 | BW25113 *dksA*::*Kan* | (2) |
| SEV161 | MG1655 *dksA*::*Kan* | This study |
| PAL2452 | MG1655 *leu82*::*Tn10 ftsZ84* | (3) |
| EAM93 | MG1655 *leu82*::*Tn10 ftsZ84* Δ*relA* | This study |
| SEA256 | MG1655 *leu82*::*Tn10 ftsZ84 dksA*::*Kan* | This study |
| MM61 | *F- araD139* Δ*lacU169 Str leu*::*Tn10 ftsA12* | (4) |
| SEA517 | MG1655 *leu82*::Tn*10 ftsA12* | This study |
| SEA522 | MG1655 *leu82*::Tn*10 ftsA12* Δ*relA* | This study |
| SEA520 | MG1655 *leu82*::Tn*10 ftsA12 dksA*::*Kan* | This study |
| WM2101 | MG1655 *lacU169 ycaD::*Tn*10 ftsK44* | (5) |
| SEA534 | MG1655 *lacU169 ycaD::*Tn*10 ftsK44* Δ*relA* | This study |
| EAM395 | MG1655 *lacU169 ycaD::*Tn*10 ftsK44 dksA*::*Kan* | This study |
| PAL2628 | MG1655 *leu82*::Tn*10 ftsQ1* | (4) |
| EAM278 | MG1655 *leu82*::Tn*10 ftsQ1 relA*::*Kan* | This study |
| EAM316 | MG1655 *leu82*::Tn*10 ftsQ1 dksA*::*Kan* | This study |
| WM4649 | MG1655 *lacU169 leu82*::Tn*10 ftsI23* | (6) |
| SEA34 | MG1655 *lacU169 leu82*::Tn*10 ftsI23 relA*::*Kan* | This study |
| EAM408 | MG1655 *lacU169 leu82*::Tn*10 ftsI23 dksA*::*Kan* | This study |
| BH330 | MG1655 $P_{lac}$-*gfp-ftsZ* | (7) |
| SEA599 | MG1655 $P_{lac}$-*gfp-ftsZ* ppGpp$^0$ | This study |
| SEA588 | MG1655 $P_{lac}$-*gfp-ftsZ dksA*::*Kan* | This study |
| EAM410 | MG1655 $P_{210}$-*gfp-ftsA* | (8) |
| SEA602 | MG1655 $P_{210}$-*gfp-ftsA* ppGpp$^0$ | This study |
| SEA592 | MG1655 $P_{210}$-*gfp-ftsA dksA*::*Kan* | This study |
| PAL3700 | MG1655 Δ*lacIZYA*::*frt* $P_{lac}$-*gfp-ftsL* | (9) |
| SEA605 | MG1655 Δ*lacIZYA*::*frt* $P_{lac}$-*gfp-ftsL* ppGpp$^0$ | This study |
| SEA589 | MG1655 $P_{lac}$-*gfp-ftsL dksA*::*Kan* | This study |
| EAM412 | MG1655 $P_{207}$-*gfp-ftsI* | (8) |
| SEA603 | MG1655 $P_{207}$-*gfp-ftsI* ppGpp$^0$ | This study |
| SEA594 | MG1655 $P_{207}$-*gfp-ftsI dksA*::*Kan* | This study |
| EAM621 | MG1655 $P_{204}$-*gfp-ftsN* | (8) |
| SEA579 | MG1655 $P_{204}$-*gfp-ftsN* ppGpp$^0$ | This study |
| SEA565 | MG1655 $P_{204}$-*gfp-ftsN dksA*::*Kan* | This study |
| RLG14538 | MG1655 $rpoZ_{\Delta 2\text{-}5}$-*kanR* $rpoC_{R362A/R417A/K615A/N680A/K681A}$-*tetAR* ($RNAP_{1\text{-}2\text{-}}$) | (10) |
| RLG14535 | MG1655 *rpoZ-kanR rpoC-tetAR* ($RNAP_{1+2+}$) | (10) |
| SEA22 | MG1655 ppGpp$^0$ *dksA*::*Kan* | This study |
| SEA640 | MG1655 $P_{lac}$-*gfp-ftsZ* ppGpp$^0$ *dksA*::*Kan* | This study |
| SEA642 | MG1655 $P_{210}$-*gfp-ftsA* ppGpp$^0$ *dksA*::*Kan* | This study |
| SEA644 | MG1655 $P_{207}$-*gfp-ftsI* ppGpp$^0$ *dksA*::*Kan* | This study |
| SEA646 | MG1655 $P_{207}$-*gfp-ftsI* ppGpp$^0$ *dksA*::*Kan* | This study |
| SEA639 | MG1655 $P_{204}$-*gfp-ftsN* ppGpp$^0$ *dksA*::*Kan* | This study |

| | | |
|---|---|---|
| CH2570 | MG1655 *lacZ*-U118 Δ*yjaZ*::*kan* Δ*btuB*3191::*Tn*10 *rpoC*$_{E677G}$ | (11) |
| SEA363 | MG1655 ppGpp$^0$ Δ*btuB*3191::*Tn*10 *rpoC*$_{E677G}$ | This study |
| BH142 | MG1655 *leu82*::*Tn10 ftsA** (*ftsA*$_{R286W}$) | (12) |
| CSW810 | MG1655 *relA*::*Kan spoT*::*cat* | This study |
| SEA385 | MG1655 *relA*::*Kan spoT*::*cat leu82*::*Tn10 ftsA** (*ftsA*$_{R286W}$) | This study |
| SEA560 | MG1655 *leuO*::*cat* pDK46 | This study |
| SEA574 | MG1655 *ftsZ84 leuO*::*cat* | This study |
| SEA609 | MG1655 *ftsZ84 leuO*::*cat rpoZ-kan rpoC-tetAR* (RNAP$_{1+2+}$) | This study |
| SEA611 | MG1655 *ftsZ84 leuO*::*cat rpoZ*$_{\Delta 2\text{-}5}$-*kanR* *rpoC*$_{R362A/R417A/K615A/N680A/K681A}$-*tetAR* (RNAP$_{1\text{-}2\text{-}}$) | This study |

### Plasmids

| Designation | Genotype | Source |
|---|---|---|
| pALS10 (*prelA*) | *lacI*$^q$ P$_{tac}$-*relA bla* | (13) |
| pALS14 (*prelA*') | *lacI*$^q$ P$_{tac}$-*relA*$_{1\text{-}331}$ *bla* | (13) |
| pRLG6332 (pINIIIA1) | *bla* | (14, 15) |
| pRLG6333 (*pdksA*) | *bla* P$_{lpp}$-P$_{lac}$-*dksA* | (14) |
| pRLG14800 (*pdksA*$_{K98A}$) | *bla* P$_{lpp}$-P$_{lac}$-*dksA*$_{K98A}$ | Gift from R. Gourse |
| pRLG14802 (*pdksA*$_{R91A}$) | *bla* P$_{lpp}$-P$_{lac}$-*dksA*$_{R91A}$ | Gift from R. Gourse |
| pRLG8874 (*pdksA*$_{D71N/D74N}$) | *bla* P$_{lpp}$-P$_{lac}$-*dksA*$_{D71N/D74N}$ | Gift from R. Gourse |
| pRLG8890 (*pdksA*$_{N88I}$) | *bla* P$_{lpp}$-P$_{lac}$-*dksA*$_{N88I}$ | Gift from R. Gourse |
| pBS58 (*pftsQAZ*) | *spc*$^R$ *ftsQAZ* | (16) |
| *pftsZ* | *spc*$^R$ *ftsZ* | This study |
| *pftsQA* | *spc*$^R$ *ftsQA* | This study |
| pCH201(*pftsN*) | *bla lacI*$_q$ *P*$_{lac}$::*gfp-FtsN* | (17, 18) |
| pKD3 | *bla frt-cat-frt* | (19) |
| pKD46 | *bla repA101ts exo bet gam araC* | (19) |

### Primers

| Designation | Use | Sequence | Source |
|---|---|---|---|
| oSEA104 | To make *leuO*::*cat* using pKD3 | GCATTCCAATAAGGGAAAGGGAGTTAA GTGTGACAGTGGAGTTAAGTATGGTGT AGGCTGGAGCTGCTTC | This study |
| oSEA105 | To make *leuO*::*cat* using pKD3 | CATTCATGTCTGACCTATTCTGCAATCA GTTAGCGTTTGCAAATTGAGACATGGGA ATTAGCCATGGTCC | This study |
| oSEA135 | To make *pftsZ* | ATGTTTGAACCAATGGAACTTAC | This study |
| oSEA136 | To make *pftsZ* | ATTAGTCCGCCAGTTCCA | This study |
| oSEA149 | To make *pftsQA* | TAAGAATTGACTGGAATTTGG | This study |
| oSEA150 | To make *pftsQA* | CATAGTTTCTCTCCGATTTG | This study |

### References

1. Guyer MS, Reed RR, Steitz JA, Low KB. 1981. Identification of a sex-factor-affinity site in E. coli as gamma delta. Cold Spring Harb Symp Quant Biol 45 Pt 1:135-40.

2. Baba T, Ara T, Hasegawa M, Takai Y, Okumura Y, Baba M, Datsenko KA, Tomita M, Wanner BL, Mori H. 2006. Construction of Escherichia coli K-12 in-frame, single-gene knockout mutants: the Keio collection. Molecular systems biology 2:2006.0008-2006.0008.
3. Arjes HA, Lai B, Emelue E, Steinbach A, Levin PA. 2015. Mutations in the bacterial cell division protein FtsZ highlight the role of GTP binding and longitudinal subunit interactions in assembly and function. BMC Microbiol 15:209.
4. Chen JC, Weiss DS, Ghigo JM, Beckwith J. 1999. Septal localization of FtsQ, an essential cell division protein in Escherichia coli. J Bacteriol 181:521-30.
5. Haeusser DP, Rowlett VW, Margolin W. 2015. A mutation in Escherichia coli ftsZ bypasses the requirement for the essential division gene zipA and confers resistance to FtsZ assembly inhibitors by stabilizing protofilament bundling. Mol Microbiol 97:988-1005.
6. Schoenemann KM, Krupka M, Rowlett VW, Distelhorst SL, Hu B, Margolin W. 2018. Gain-of-function variants of FtsA form diverse oligomeric structures on lipids and enhance FtsZ protofilament bundling. Mol Microbiol 109:676-693.
7. Hill NS, Buske PJ, Shi Y, Levin PA. 2013. A Moonlighting Enzyme Links Escherichia coli Cell Size with Central Metabolism. PLOS Genetics 9:e1003663.
8. Mueller EA, Westfall CS, Levin PA. 2020. pH-dependent activation of cytokinesis modulates Escherichia coli cell size. PLoS Genet 16:e1008685.
9. Tsang MJ, Bernhardt TG. 2015. A role for the FtsQLB complex in cytokinetic ring activation revealed by an ftsL allele that accelerates division. Mol Microbiol 95:925-44.
10. Ross W, Sanchez-Vazquez P, Chen AY, Lee JH, Burgos HL, Gourse RL. 2016. ppGpp Binding to a Site at the RNAP-DksA Interface Accounts for Its Dramatic Effects on Transcription Initiation during the Stringent Response. Mol Cell 62:811-823.
11. Satory D, Halliday JA, Sivaramakrishnan P, Lua RC, Herman C. 2013. Characterization of a novel RNA polymerase mutant that alters DksA activity. J Bacteriol 195:4187-94.
12. Hill NS, Kadoya R, Chattoraj DK, Levin PA. 2012. Cell size and the initiation of DNA replication in bacteria. PLoS Genet 8:e1002549.
13. Svitil AL, Cashel M, Zyskind JW. 1993. Guanosine tetraphosphate inhibits protein synthesis in vivo. A possible protective mechanism for starvation stress in Escherichia coli. J Biol Chem 268:2307-11.
14. Rutherford ST, Lemke JJ, Vrentas CE, Gaal T, Ross W, Gourse RL. 2007. Effects of DksA, GreA, and GreB on transcription initiation: insights into the mechanisms of factors that bind in the secondary channel of RNA polymerase. J Mol Biol 366:1243-57.
15. Masui Y, Mizuno T, Inouye M. 1984. Novel High-level Expression Cloning Vehicles: 104-fold Amplification of Escherichia coli Minor Protein. Bio/Technology 2:81-85.
16. Bi E, Lutkenhaus J. 1990. FtsZ regulates frequency of cell division in Escherichia coli. Journal of Bacteriology 172:2765-2768.
17. Gerding MA, Liu B, Bendezú FO, Hale CA, Bernhardt TG, de Boer PA. 2009. Self-enhanced accumulation of FtsN at Division Sites and Roles for Other Proteins with a SPOR domain (DamX, DedD, and RlpA) in Escherichia coli cell constriction. J Bacteriol 191:7383-401.
18. Hale CA, de Boer PA. 2002. ZipA is required for recruitment of FtsK, FtsQ, FtsL, and FtsN to the septal ring in Escherichia coli. J Bacteriol 184:2552-6.
19. Datsenko KA, Wanner BL. 2000. One-step inactivation of chromosomal genes in Escherichia coli K-12 using PCR products. Proceedings of the National Academy of Sciences of the United States of America 97:6640-6645.
